# Interactions of Sleep and Repetitive Behavior in Attention-Deficit/Hyperactivity Disorder, Autism and Intellectual Disability

**DOI:** 10.64898/2025.12.04.692304

**Authors:** Maria Niedernhuber, Giorgia Michelini, Margherita Malanchini, Afia Ali, Valdas Noreika

**Author notes:** Correspondence concerning this article should be addressed to Maria Niedernhuber, School of Biological and Behavioural Sciences, Queen Mary University of London, Mile End Road, London E1 4NS. Author note: The authors made the following contributions. Maria Niedernhuber: Conceptualization, Writing - Original Draft Preparation, Writing - Review & Editing, Formal analysis, Visualization; Giorgia Michelini: Writing - Review & Editing, Supervision; Margherita Malanchini: Writing - Review & Editing, Supervision; Afia Ali: Writing - Review & Editing, Supervision; Valdas Noreika: Writing - Review & Editing, Supervision, Project administration.

## Abstract

**Objective:** Sleep problems and restricted and repetitive behaviors (RRBs) feature in neurodevelopmental conditions such as autism, attention-deficit/hyperactivity disorder (ADHD), and intellectual disabilities (ID). It is unclear whether their interaction differs between conditions, and whether symptom dimensions might capture differences better than diagnostic categories. Method: In a preregistered cross-sectional analysis of the Healthy Brain Network community sample enriched for neurodevelopmental conditions (n = 2,394, 5–21 years), linear regression modeled parent-reported sleep disturbance and RRBs by diagnostic category and symptom dimensions (autistic traits, ADHD traits, IQ)

**Results:** All symptom dimensions predicted sleep disturbance (p < .001), whereas only ADHD consistently predicted greater sleep disturbance (d = .35, 95% CI [.24, .47]) among diagnostic categories relative to typical development. All diagnostic categories and symptom dimensions were associated with RRB severity. Sleep disturbance and RRBs were associated in the entire sample (β = .30, p < .001). Sleep disturbance predicted RRBs at all levels of autistic and ADHD traits, but most strongly with high autistic and low ADHD traits (β = .22) and least strongly when both were high (β = .13). Dimensional measures explained more variance than diagnoses for sleep disturbance (R² = .14 vs .03) and RRBs (R² = .38 vs .21), and provided better model fits in all comparisons.

**Conclusion:** Autistic and ADHD traits shaped the relationship between sleep disturbance and RRBs, whereas diagnoses did not. Neurodevelopmental symptom assessments might be more informative than diagnoses to manage sleep and RRBs, though the cross-sectional design and parent report limit inference.

**Plain language summary:** Youth with neurodevelopmental conditions often sleep badly. We asked how sleep relates to repetitive behavior and narrow interests, and if this differs between autism, ADHD, and intellectual disability. We surveyed caregivers of 2,400 young people aged 5–21 referred for mental health or learning concerns. Poorer sleep accompanied repetitive behaviors regardless of diagnosis. The link was strongest with more autistic and fewer ADHD symptoms. Symptoms might guide support better than diagnoses.

## Introduction

Sleep problems are common health concerns disproportionately affecting individuals with neurodevelopmental conditions^1^. Examples of neurodevelopmental conditions with higher rates of sleep disturbance include autism, Attention-Deficit/Hyperactivity Disorder (ADHD), and intellectual disabilities (ID). Common sleep issues in autism include prolonged sleep latency, frequent awakenings at night, and reduced sleep duration^1^, and this is linked to emotional, cognitive, behavioral and social difficulties^2^. Children with ADHD often experience sleep issues such as delayed sleep onset, and insomnia^1^, which are linked to impaired executive function, adaptive behavior, low academic achievement and mental health issues^3^. There is mixed evidence whether stimulant medication might influence sleep disturbance in ADHD^4^. Likewise, people with ID spend fewer hours asleep, and experience poorer sleep quality than typically developing individuals^5^. This subgroup also exhibits challenging behavior, increased symptom severity, and rates of neurological and psychiatric comorbidity^6^.

Sleep, cognition and sensory processing are intimately related, and crucial for development^7^. Accordingly, sleep difficulties feature prominently in neurodevelopmental conditions, which are characterized by altered sensorimotor^8,9^ and cognitive function^10^. Atypical development can manifest as restricted and repetitive behaviors (RRBs) which encompass both altered sensorimotor processing (such as sensation-seeking, and over- or underreactivity to sensory input, stereotyped motor movements, and atypical object manipulation) and cognitive patterns (such as repetitive speech, strict daily routines, intense distress in response to change, and restricted interests)^11^. Such behaviors are a core characteristic of autism and also occur in ADHD and ID, albeit not to the same extent^12,13^. For instance, sensation-seeking behavior features in autism^14^, ADHD^8^, and ID^9^. Likewise, reduced cognitive flexibility was observed in ADHD^15^, autism^16^, and ID^17^.

Multiple studies support a link between sleep dysfunction and RRBs in autistic children and young people^18^. Although fewer studies address this interaction in other neurodevelopmental conditions, there is also some evidence for a relationship between sleep dysfunction and RRBs in children with developmental delay^18^ and ADHD^19^. For instance, children with developmental delay who exhibit higher-order RRBs also reported more sleep disturbance^18^. In people with ID, sleep disturbance was associated with stereotyped behavior^20^. Similarly, children and adolescents with ADHD who have sleep problems also experience issues with sensory over- and underresponsivity^19^.

An open question is whether there are systematic differences in the interaction between sleep difficulties and RRBs between neurodevelopmental conditions. There is evidence that both RRBs and sleep dysfunction manifest differently across the three conditions. For instance, RRBs feature more prominently in autism than ADHD or ID^12,21^. Daytime sleepiness is more prominent in ADHD than autism^22^. Bedtime resistance is more prominent in ID than autism, whereas insomnia is more common in the latter^23^. Only pairwise comparisons between conditions flag differences in sleep dysfunction and RRBs, and a comparison of all three conditions is still lacking. Due to substantial symptom overlap between neurodevelopmental conditions, as well as high rates of co-occurring diagnoses^24^, it is challenging to determine differences in the interaction between sleep and RRBs between neurodevelopmental conditions. Previous studies found that rates of both sleep disturbance and RRBs increase when multiple diagnoses are present. For example, in autistic children, co-occurring ID was found to contribute to sensation-seeking (e.g. self-stimulation with loud sounds)^13^, as well as ritualistic and repetitive behaviors^25^. Autistic children with ADHD report more sleep disturbance and poorer sleep hygiene^26^. In people with ID, RRBs were associated with the ADHD traits hyperactivity and impulsivity^27^.

To address this issue, it might be more useful to apply a transdiagnostic approach focusing on neurodevelopmental dimensional measures rather than binary diagnostic categories. An alternative framework termed the neurodevelopmental spectrum describes individuals using transdiagnostic dimensional measures shared between different conditions rather than distinct conditions^24,28^. Autism and ADHD are common co-existing conditions among young people. 37–78% of autistic children also have ADHD^29^, and ∼20-50% of children with ADHD also qualify for an autism diagnosis^30^. Even when there is no dual diagnosis, children with ADHD and autism present with similar difficulties, including impairments in social function and RRBs^12^. Therefore, we suggest that using both symptom dimensions and diagnostic categories will provide a more comprehensive view of the relationship between sleep disturbance and RRBs in neurodevelopmental conditions.

In a preregistered analysis (AsPredicted #296353), we systematically examined the cross-sectional relationship of sleep disturbance and RRBs with autism, ADHD, and ID, as well as tested whether the relationship between sleep and RRBs is moderated by these neurodevelopmental conditions. First, we hypothesized that individuals in the neurodevelopmental groups would exhibit significantly greater sleep disturbance than controls (H1). Second, we predicted RRBs to feature most prominently in autism, followed by ADHD and ID, and finally TD (H2). Third, we expected a cross-sectional relationship between sleep and RRBs across all participants (H3). Since RRBs are most extensively observed in autism^12^, we reasoned that this relationship would be strongest in this group, followed by ADHD, ID and weakest in TD (H4). All hypotheses were tested based on (a) diagnostic categories and (b) dimensional measures of neurodevelopmental traits. Due to the clinical overlap between ADHD and autism^29,30^, we also investigated whether the co-occurrence of autism with either of these conditions might compound the influence of autism alone on sleep disturbance, RRBs, or their interaction. Therefore, we tested for synergistic or additive effects of concurrent autism and ADHD, as well as autism and ID whenever neurodevelopmental conditions were compared.

## Methods

### Participants

We performed a secondary data analysis of the Healthy Brain Network (HBN) project sample, which is a large-scale, open-access study initiative focused on child and adolescent mental health^31^. The HBN project recruited participants aged 5–21 seeking out families concerned about their child/children’s mental health and learning, using a community-referred sampling model from different community settings located in New York, USA (e.g., schools, clinics, community events). As a result, this sample includes higher rates of neurodevelopmental conditions than expected in the general population^31^. More than one child per family could be recruited. Data until the latest data release (#11, n = 4867) were used in the current analysis. In our study, participants were assigned to one or multiple of the following conditions: ADHD, autism, ID, and TD. Assignment to the autism and ADHD groups relied on clinician consensus diagnoses. Specifically, inclusion in the autism group required an ICD-10 diagnosis of F84. Individuals in the ADHD group required an ICD-10 diagnosis of F90 (F90.0, F90.1, F90.2, F90.8, F90.9). Participants completed a web-based version of the Kiddie Schedule for Affective Disorders and Schizophrenia (K-SADS), which is a semi-structured interview examining current and past evidence of psychiatric conditions in children and adolescents based on the DSM-5^32^. Based on clinical interviews and reviews of study participation materials, the clinical team generated consensus diagnoses which were used in our study.

For the purpose of this study, participants had to meet two criteria to be categorized as having ID: (1) a Full-Scale IQ (FSIQ) score of less than 70; (2) a standard score below 85 on at least one domain from the VABS (Communication, Daily Living Skills, or Socialization), measuring adaptive functioning. We did not classify 34 participants with an IQ < 70 (mean IQ = 63.0, SD = 7.22, range = 40-69) as having ID. One participant did not meet the cutoff in the Vineland Adaptive Behavior Scale (VABS; Communication = 93, Socialization = 85, Daily Living Skills = 85). For the remaining participants, VABS data was incomplete. Participants without any neurological/psychiatric diagnoses (ICD-10 F- or G-code) were categorized as TD.

In total, 3,333 participants had complete data on age, sex, BSMSS, IQ, SDSC and RBS-R. We excluded 197 because no clinician consensus diagnosis was recorded. A further 742 participants who did not meet criteria for inclusion in either the autism, ADHD, ID group also could not be classified as TD since they were diagnosed with a neurological or psychiatric condition (ICD-10 F or G). This resulted in a sample of 2394 participants. Based on the ICD-10, autistic people were diagnosed with F84.0 (Childhood Autism Spectrum Disorder). 861 (44.2%) individuals with ADHD were predominantly inattentive (F90.0), 107 (5.5%) were predominantly hyperactive (F90.1), 853 (43.8%) presented with a combined type (F90.2), 125 (6.4%) with other type (F90.8), and 2 (0.1%) with an unspecified type (F90.9). Figure 1 A provides an overview of concurrent conditions.

**Figure 1.**
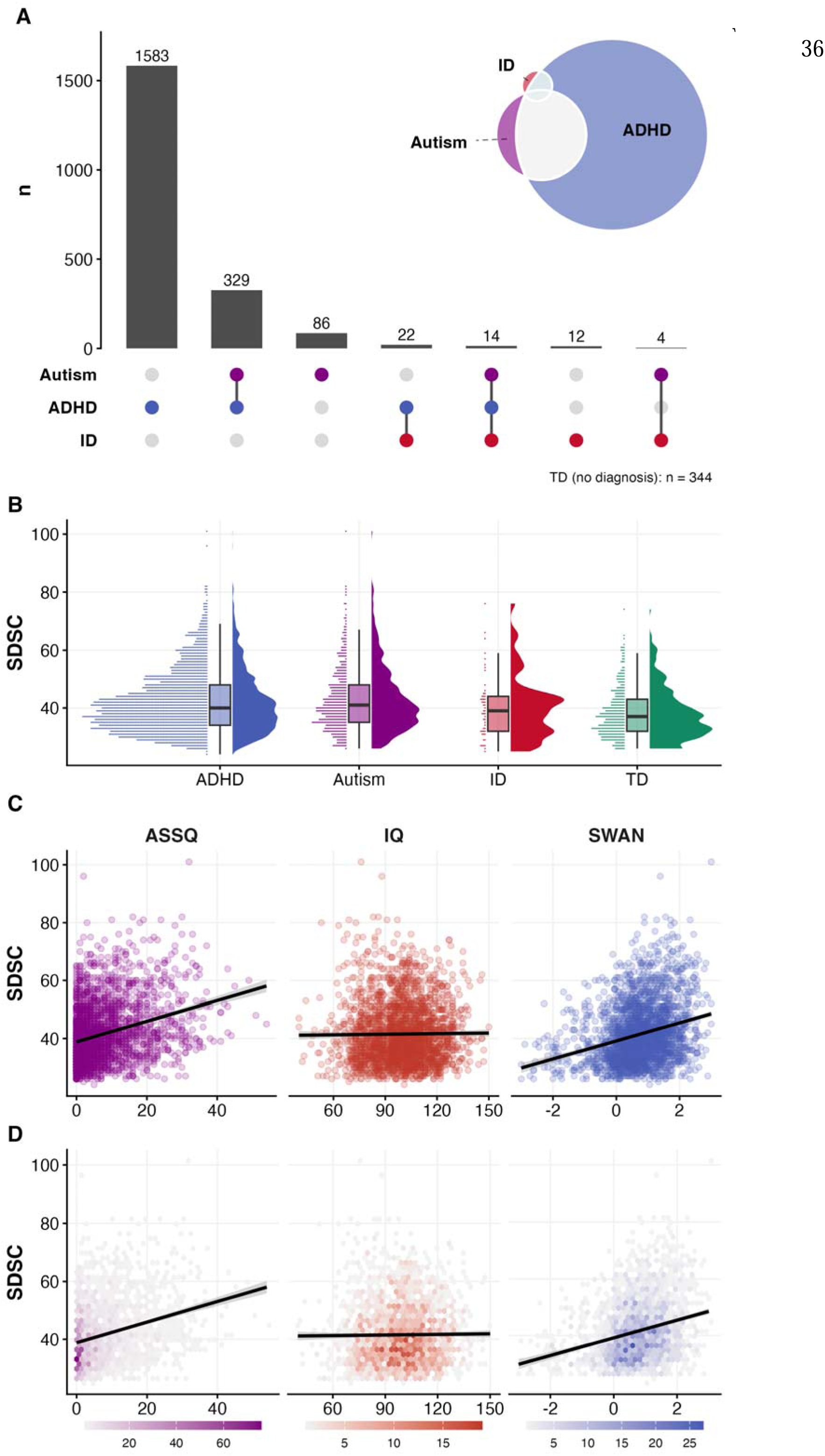
(A) Co-occurrence of autism, ADHD and ID. (B) Distribution of SDSC scores across diagnostic groups. Raincloud plots display probability density, boxplots (median and interquartile range), and a scatterplot showing individual ratings. (C) Relationship between SDSC score and symptom dimensions for autism (ASSQ), ID (IQ), and ADHD (SWAN) represented with a linear regression line, and shaded regions representing 95% confidence intervals. (D) Bivariate density distributions (hexagonal binning) mapping ASSQ, IQ, and SWAN scores to SDSC, overlaid with linear regression lines. Color intensity indicates the observation count within each bin.

In our ID group, 41 (78.8%) of 52 were diagnosed with intellectual disability (F70 mild, n = 35; F71 moderate, n = 6). One participant received a diagnosis of borderline intellectual functioning (R41.83). Co-occurring diagnoses in this group most commonly included ADHD (n = 36, 69.2%), autism (n = 18, 34.6%), developmental speech and language disorder (n = 12, 23.1%), and specific learning disorder with impairment in reading (n = 12, 23.1%). An overview of diagnoses per diagnostic category is shown in Supplementary Table 1.

Our sample was aged on average 9.99 ± 3.02 years (range 5.06-21.9). Guardians reported demographic information, including race and ethnicity (Table 1). Caregivers reported the sex assigned at birth, recorded as male or female. 69.4% were reported as male and 30.6% female. 54.9% reported the participants’ race as White, 13.2% as Black/African American, 9.1% as Hispanic, 2.9% as Asian, and 16.6% as of two or more races (1.6% missing). 23.6% reported the participants’ ethnicity as Hispanic/Latino. 54.5% of guardians reported a graduate degree as their highest education, 30.7% a bachelor’s degree, 9.2% some college, and 5.6% high school or less. 72.2% of guardians were married or in a domestic partnership, and 14.0% were separated or divorced (1.6% missing). Socioeconomic status, indexed by the BSMSS, averaged 50.23 ± 13.96, and FSIQ averaged 99.29 ± 17.00. Ethical approval for the original data was obtained by the Chesapeake Institutional Review Board. Participants’ legal guardians gave written consent for their child to take part in the study, and children provided written assent^31^. We did not require further ethical approval for the re-analysis of anonymized open-source data.

**Table 1:**
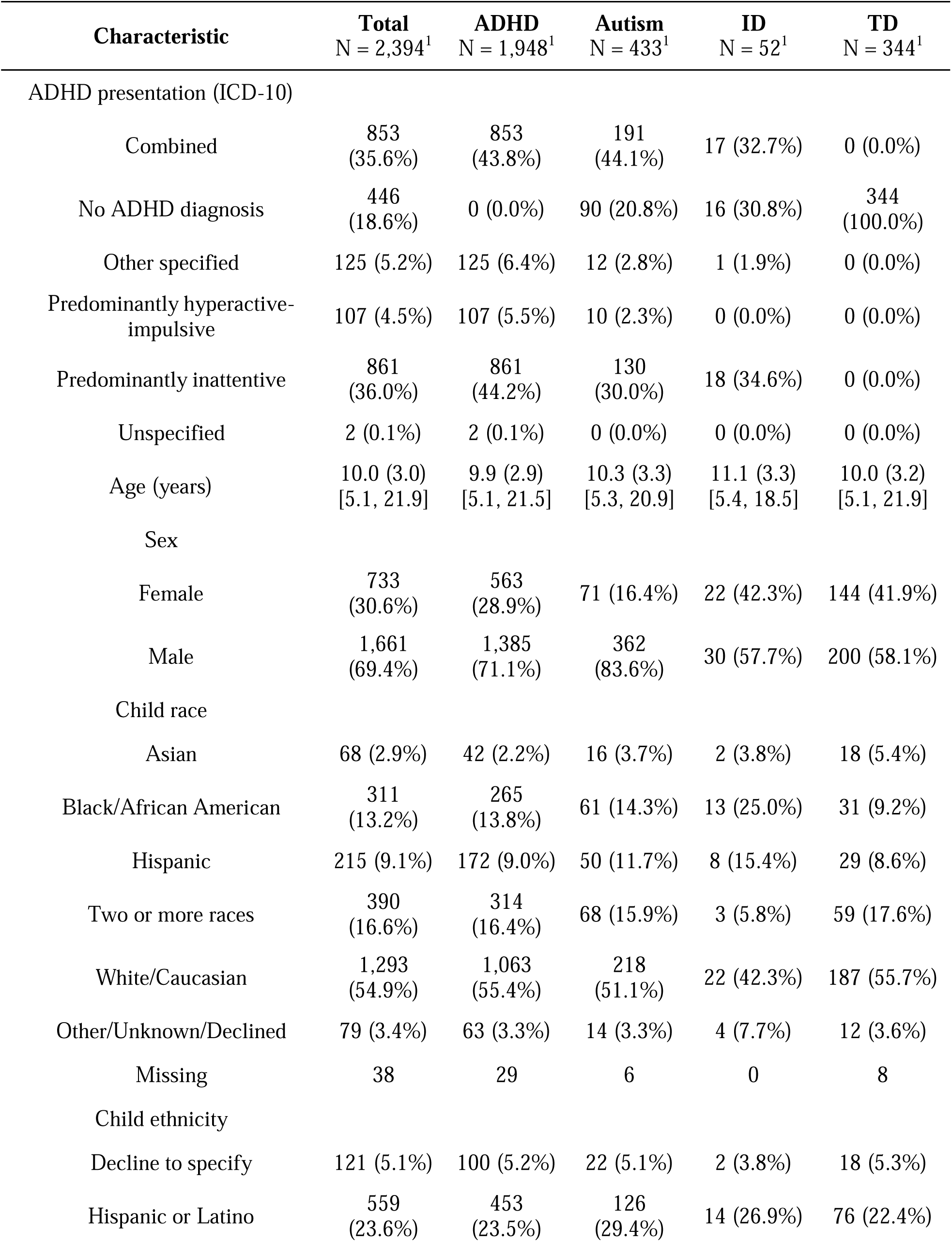
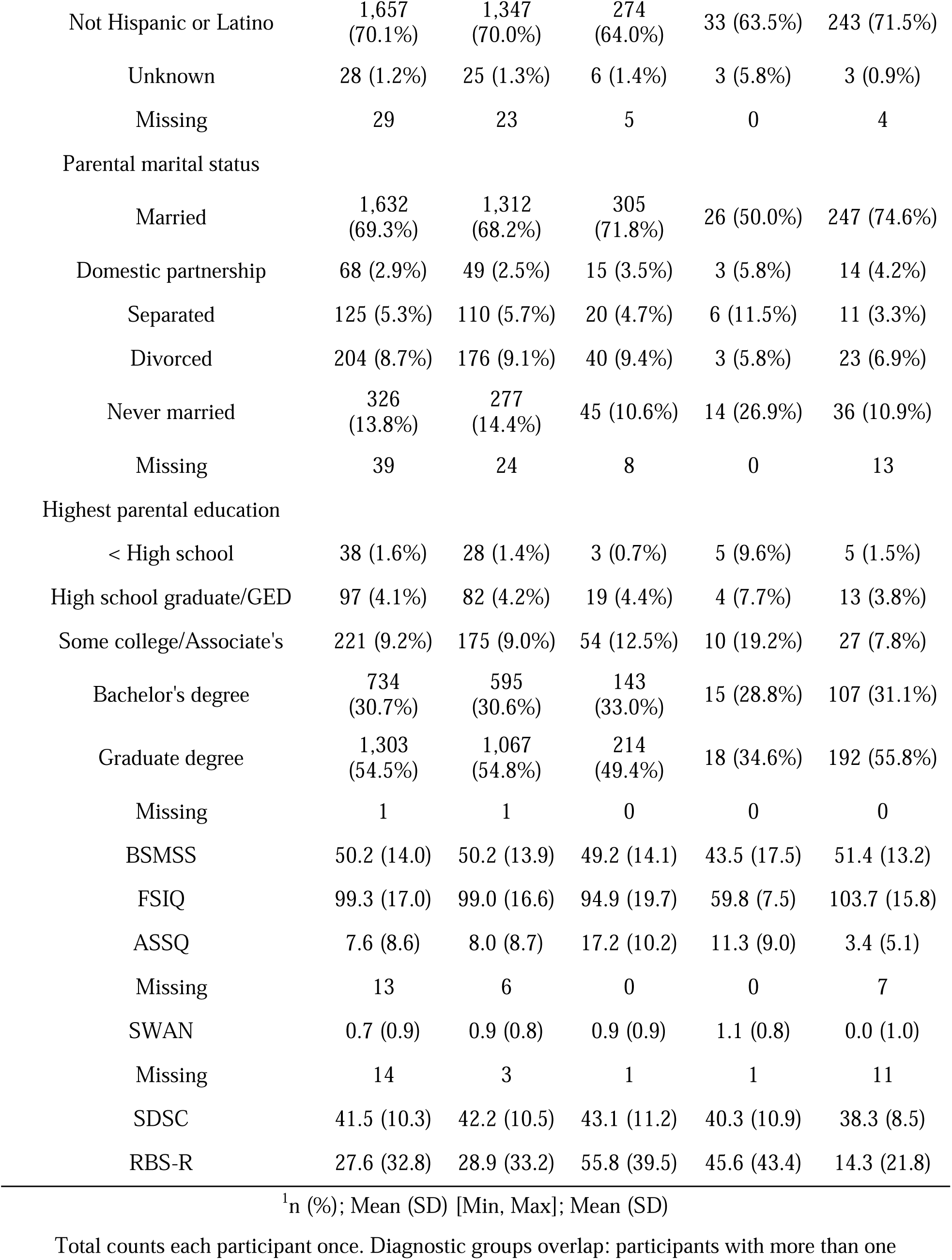
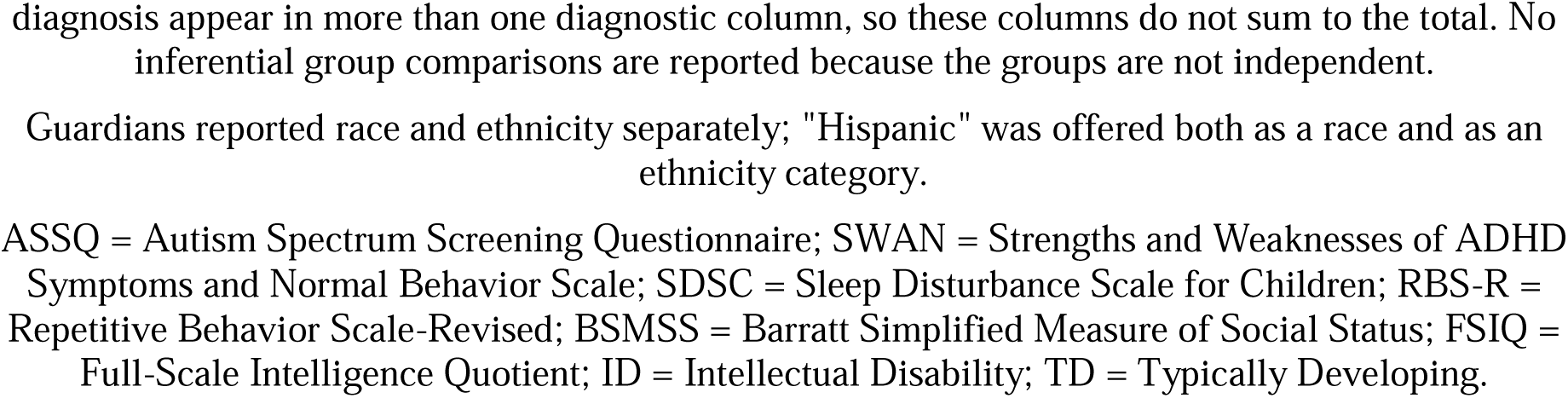
Demographic information and descriptive statistics of scale measurements.

### Measures

We measured sleep disturbance using the Sleep Disturbance Scale for Children (SDSC;^33^), and RRBs with the Repetitive Behavior Scale-Revised (RBS-R;^11^). Autistic traits were quantified using the Autism Spectrum Screening Questionnaire (ASSQ;^34^) and ADHD traits using the Strengths and Weaknesses of ADHD Symptoms and Normal Behavior Scale (SWAN;^35^). FSIQ (M = 100, SD = 15) was measured using the Wechsler Adult Intelligence Scale-IV (WAIS-IV) for participants older than 17^36^, the Wechsler Intelligence Scale for Children-V (WISC-V) for participants typically aged 6-17^37^, and the Kaufman Brief Intelligence Test-II (KBIT-2) for participants under 6 years^38^. Socioeconomic status was measured using the parent-reported Barratt Simplified Measure of Social Status (BSMSS)^39^, and adaptive functioning using the Vineland Adaptive Behavior Scales (VABS)^40^. The VABS was administered only where the clinical team identified low cognitive ability. Caregivers were asked what sex a child was assigned at birth. Sex was recorded as a binary variable (male/female). All scales were used dimensionally. Supplementary Table 2 details all measures.

### Analysis

We performed a series of preregistered analyses (AsPredicted #296353) to understand the interaction of sleep disturbance (SDSC) and RRBs (RBS-R) in autism, ADHD and ID in R version 4.6.1 (2026-06-24) using RStudio (v2026.05.1+225). Hypotheses were evaluated using diagnostic categories in the ICD-10 as well as using trait dimensions associated with each condition, using the ASSQ for autistic traits, the SWAN for ADHD traits, and IQ for ID. To test our hypotheses, we used linear regression models with age, sex, and BSMSS as covariates. SDSC, RBS-R, ASSQ, SWAN and IQ were mean-centered before modeling. Our preregistration included linear mixed-effects models with family identification number as a random intercept. However, only one family contributed more than one participant with complete data on all variables included in the model, and therefore, we fitted models with ordinary least squares. Before refitting the model to obtain standardized estimates, we z-scored the outcome variable and all continuous predictors. As a result, we used standardized β coefficients as estimates for continuous predictors, and Cohen’s d for diagnostic categories. To control for multiple comparisons, we applied FDR-correction using the Benjamini–Hochberg procedure only to key model terms for each hypothesis family. We did not FDR-correct p-values for the covariates. To run post-hoc analyses for interaction effects for models with continuous moderators, we used simple slopes analyses (interactions package v1.2.0) and tested each moderator at the 16th, 50th and 84th percentiles of its observed distribution. For categorical models, we estimated marginal means and trends (emmeans package v1.11.0). We used Tukey-adjustments for pairwise contrasts between diagnostic categories, whereas simple slopes were reported without adjustment. We excluded participants with incomplete data in SDSC, RBS-R, Age, Sex, BSMSS, IQ for all analyses. For analyses including other scales, we only removed participants with incomplete data specific to the survey from the dataset. We set the TD group as the reference level for all models.

### Exploratory analyses

We compared dimensional and diagnostic model fits using Vuong tests, and estimated R² differences with 95% percentile bootstrap confidence intervals (2,000 runs). Linear regression models adjusted for age, sex, BSMSS, and SWAN tested whether stimulant medication predicted greater sleep disturbance or higher RRB severity (FDR-corrected) in a subsample with ADHD and available caregiver-reported current psychiatric medication data (n = 1,783, stimulant: n = 390, non-stimulant n = 28, other psychiatric medication only n = 24, no psychiatric medication (n = 1,341)). To test whether RRB severity is highest in autism followed by ADHD/ID and then TD, we fitted models encompassing all diagnostic categories and covariates, referenced against the TD group, as well as a model with autistic and ADHD traits as well as reversed FSIQ. Per model, coefficients were tested against zero and compared with each other using general linear hypothesis tests (Holm-corrected).

## Results

### Sleep disturbances are linked to ADHD diagnosis and broadly associated with autistic and ADHD traits as well as IQ

Categorically, ADHD predicted more sleep disturbance than TD (*d* = .35, 95% CI [.24, .47], *t* = 6.08, *p* < .001), whereas ID (*d* = -.17, 95% CI [-.45, .10], *t* = −1.25, *p* = .281) and autism (*d* = .22, 95% CI [-.01, .45], *t* = 1.85, *p* = .102) did not. Autism predicted more sleep disturbance when modeling its interaction with ID (*d* = .18, 95% CI [.08, .29], *t* = 3.38, *p* = .001) but not ADHD (*d* = .22, 95% CI [-.01, .45], *t* = 1.85, *p* = .102). Interactions between autism and ADHD (*d* = -.03, 95% CI [-.29, .23], *t* = -.23, *p* = .816) or autism and ID (*d* = .29, 95% CI [-.28, .86], *t* = .99, *p* = .397) were not associated with sleep disturbance.

Dimensionally, higher ADHD traits (*β* = .22, 95% CI [.18, .26], *t* = 10.82, *p* < .001), higher autistic traits (*β* = .24, 95% CI [.19, .28], *t* = 9.92, *p* < .001), and higher IQ (*β* = .10, 95% CI [.06, .14], *t* = 5.11, *p* < .001) each independently predicted more sleep disturbance. There was neither an interaction of autistic and ADHD traits (*β* = .01, 95% CI [-.02, .05], *t* = .65, *p* = .547) nor autistic traits and IQ (*β* = -.02, 95% CI [-.05, .02], *t* = -.88, *p* = .432), which replicates findings from categorical models. Results for H1 can be found in Figure 1.

### RRBs are associated with interacting ADHD and autistic traits

Categorically, autism (*d* = 1.13, 95% CI [.92, 1.34], *t* = 10.69, *p* < .001), ADHD (*d* = .25, 95% CI [.14, .35], *t* = 4.69, *p* < .001), and ID (*d* = .41, 95% CI [.16, .65], *t* = 3.21, *p* = .002) each independently predicted higher RRB severity. We did not find evidence supporting a synergistic effect of autism and ADHD on RRBs (*d* = -.14, 95% CI [-.38, .09], *t* = −1.22, *p* = .221), which is consistent with an additive effect.

Dimensionally, higher autistic traits (*β* = .52, 95% CI [.48, .55], *t* = 25.36, *p* < .001), higher ADHD traits (*β* = .11, 95% CI [.08, .14], *t* = 6.04, *p* < .001), and lower IQ (*β* = -.07, 95% CI [-.10, -.03], *t* = −3.77, *p* < .001) all predicted greater RRB severity. Unlike in our categorical model, there was an interaction of autistic and ADHD traits (*β* = .04, 95% CI [.01, .07], *t* = 2.47, *p* = .017). Autistic traits predicted greater RRB severity at all levels of ADHD traits, but this effect strengthened as ADHD traits increased (16th percentile of SWAN: *β* = .48, 95% CI [.43, .53], *t* = 18.38, *p* < .001; 84th percentile: *β* = .55, 95% CI [.51, .59], *t* = 28.88, *p* < .001).

Categorically, autism interacted with ID (*d* = .54, 95% CI [.02, 1.06], *t* = 2.04, *p* = .048). There was a stronger link between autism and RRBs in individuals with ID (*d* = 1.54, 95% CI [.87, 2.21], *t* = 5.90, *p* < .001) than without (*d* = 1.00, 95% CI [.87, 1.12], *t* = 20.39, *p* < .001). Dimensionally, there was a negative interaction between autistic traits and IQ (*β* = -.06, 95% CI [-.09, -.03], *t* = −3.79, *p* < .001). As IQ increased, the relationship between autistic traits and RRBs attenuated (16th percentile of IQ: *β* = .58, 95% CI [.54, .63], *t* = 26.53, *p* < .001; 84th percentile: *β* = .46, 95% CI [.42, .51], *t* = 18.80, *p* < .001). Results are found in Figure 2.

**Figure 2.**
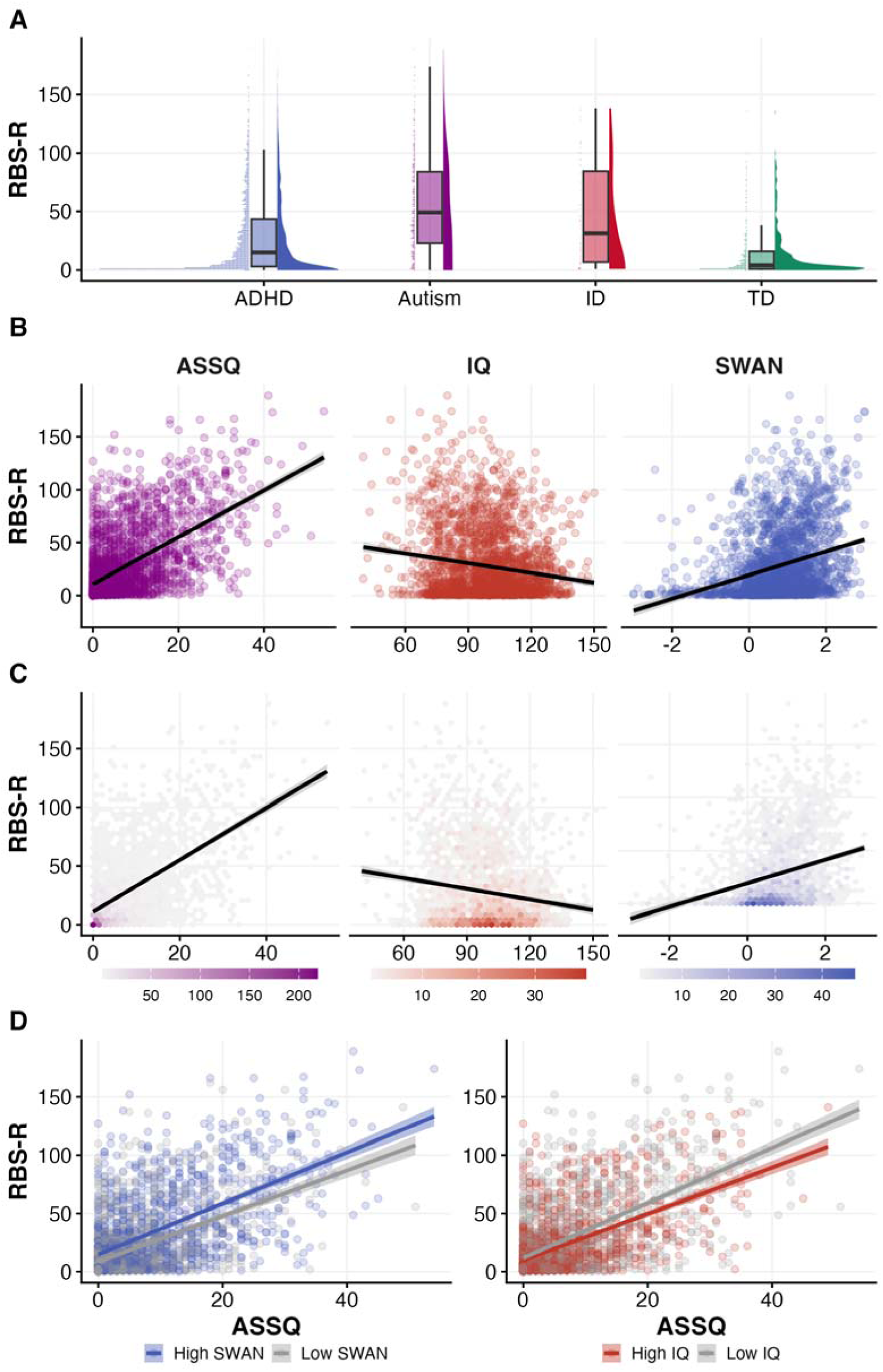
(A) Distribution of RBS-R scores across diagnostic groups, each visualized using a probability distribution slab, a boxplot with median and interquartile range, and a scatterplot. (B) Linear associations between dimensional trait measures (ASSQ, IQ, SWAN) and RBS-R, including 95% confidence intervals shown as shaded regions. (C) Hexagonal bin plots illustrating the density of observations for each trait against RBS-R, overlaid with linear regression lines. (D) The relationship between ASSQ and RBS-R, stratified by SWAN (left) and IQ (right) using a median split. Regression lines are shown in blue/grey for high/low SWAN (left) and in red/grey for high/low IQ to illustrate moderating effects.

### Neurodevelopmental traits, but not categorical diagnoses, moderate the relationship between sleep disturbance and RRBs

More sleep disturbance was associated with higher RRB severity across the whole sample (*β* = .30, 95% CI [.27, .34], *t* = 15.91, *p* < .001), and vice versa (*β* = .32, 95% CI [.28, .35], *t* = 15.91, *p* < .001; Figure 3A).

**Figure 3.**
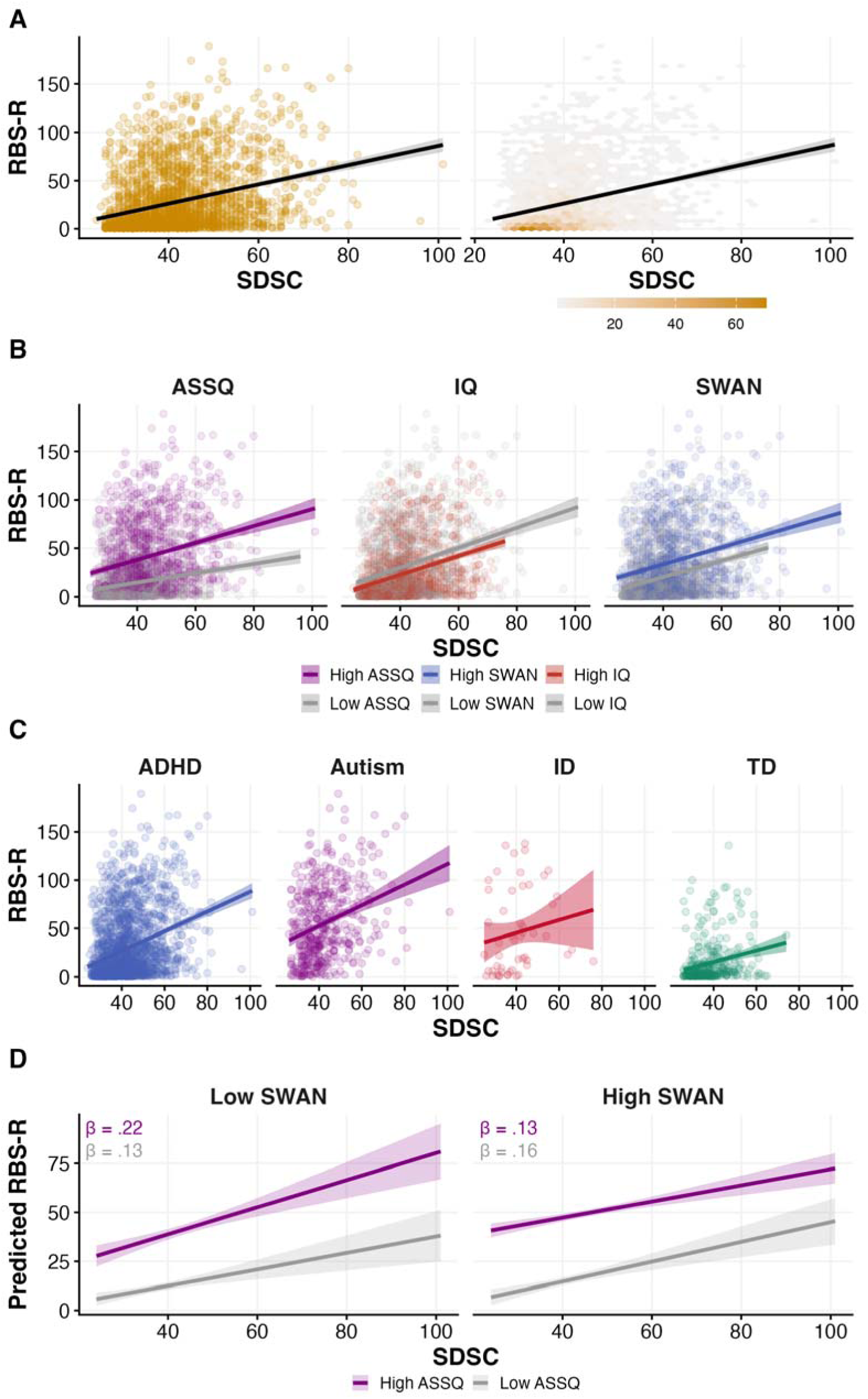
(A) Relationship between SDSC and RBS-R across the entire sample, represented as a standard scatterplot adjacent to a hexagonal density plot with a regression line and shaded 95% confidence intervals. (B) Relationship between SDSC and RBS-R, faceted by dimensional symptom measure (ASSQ, IQ, SWAN). Each facet shows high and low trait levels split along the median, using regression lines (with shaded 95% confidence intervals). (C) Relationship between SDSC and RBS-R shown using scatterplots and regression lines (with shaded 95% confidence intervals) for each diagnostic group. (D) Simple slopes represent SDSC predicting RBS-R from the three-way interaction between SDSC, ASSQ and SWAN. Lines highlight predicted RBS-R for low (16th percentile) and high (84th percentile) ASSQ, faceted by low (16th percentile) and high SWAN (84th percentile), adjusted for age, sex, BSMSS and IQ, with shaded 95% confidence intervals. Labels give standardized simple slopes (β).

Categorically, sleep disturbance (*β* = .19, 95% CI [.09, .30], *t* = 3.59, *p* < .001) predicted higher RRBs, and vice versa (*β* = .29, 95% CI [.14, .43], *t* = 3.89, *p* < .001). ADHD (*d* = .18, 95% CI [.07, .28], *t* = 3.28, *p* = .002), autism (*d* = 1.09, 95% CI [.88, 1.29], *t* = 10.67, *p* < .001), and ID (*d* = .44, 95% CI [.20, .68], *t* = 3.66, *p* < .001) were positively associated with RRBs. Crucially, no three-way interaction involving diagnostic categories reached significance (all *p* > .05). Co-occurring autism and ID were associated with higher RRB severity (*d* = .58, 95% CI [.08, 1.09], *t* = 2.36, *p* = .035).

Unlike categorical models, dimensional models revealed a three-way interaction between sleep disturbance, autistic traits, and ADHD traits predicting RRBs (*β* = -.04, 95% CI [-.06, -.01], *t* = −2.47, *p* = .027). In a simple slopes analysis, sleep disturbance predicted RRBs at every combination of traits. We found that the association was steepest when autistic traits were high and ADHD traits low (*β* = .22, 95% CI [.14, .29], *t* = 5.58, *p* < .001) but shallowest when both were high (*β* = .13, 95% CI [.09, .17], *t* = 5.84, *p* < .001). However, the equivalent three-way interaction predicting sleep disturbance from RRBs was not significant (*β* = -.01, 95% CI [-.04, .01], *t* = -.89, *p* = .427). We found that the relationship between RRBs and sleep disturbance was moderated by autistic traits (*β* = -.05, 95% CI [-.08, -.01], *t* = −2.38, *p* = .034). The link between RRBs and sleep disturbance attenuated as autistic traits increased (16th percentile: *β* = .27, 95% CI [.21, .34], *t* = 8.19, *p* < .001; 84th percentile: *β* = .18, 95% CI [.13, .24], *t* = 6.75, *p* < .001). IQ also moderated this association (*β* = .07, 95% CI [.02, .11], *t* = 2.90, *p* = .008). Higher RRB severity predicted more sleep disturbance more strongly as IQ increased (16th percentile: *β* = .17, 95% CI [.11, .23], *t* = 5.64, *p* < .001; 84th percentile: *β* = .30, 95% CI [.23, .38], *t* = 8.08, *p* < .001). Results for H4 are shown in Figure 3B–D. A comprehensive overview of our results is provided in Table 2.

**Table 2:**
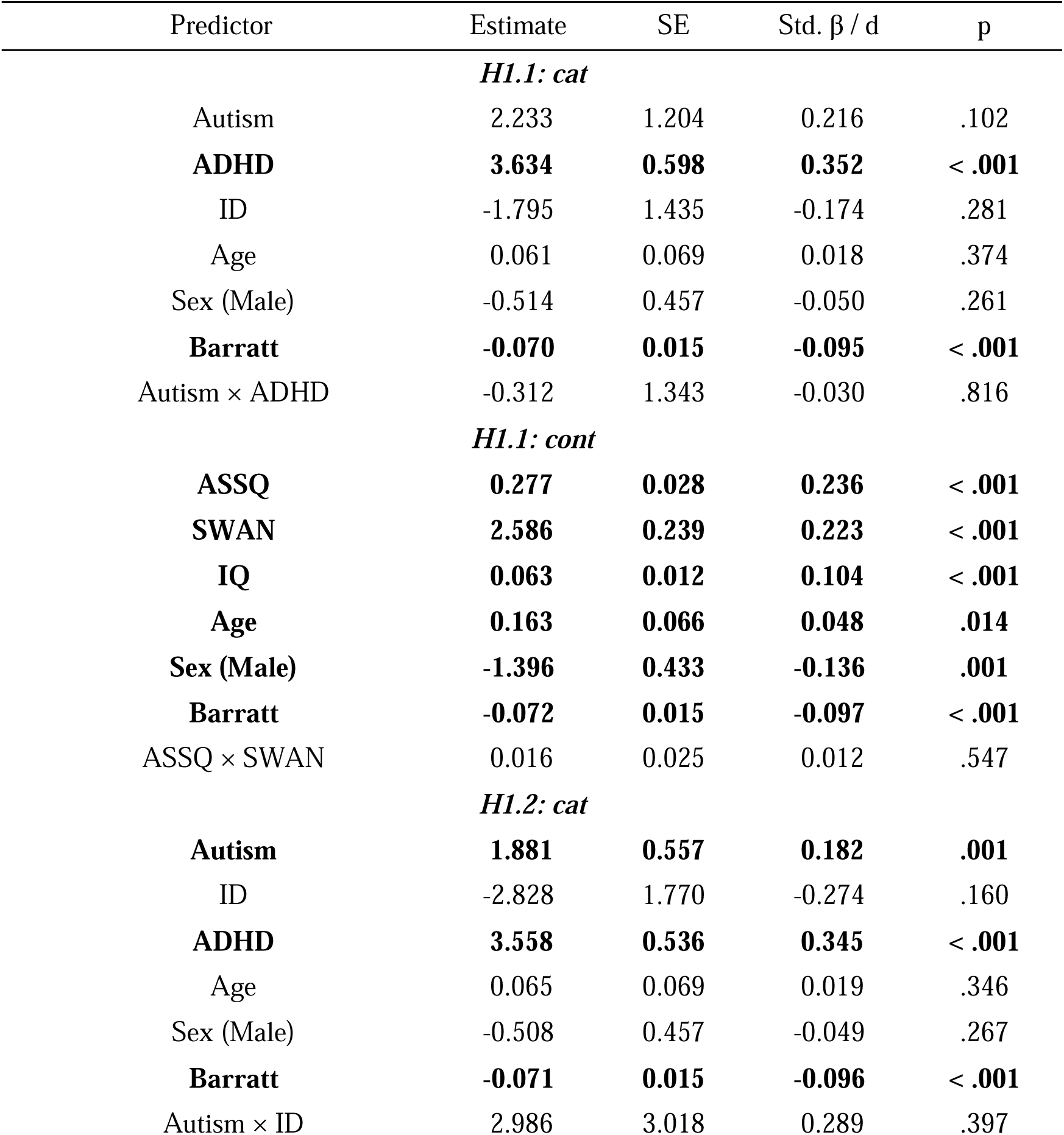
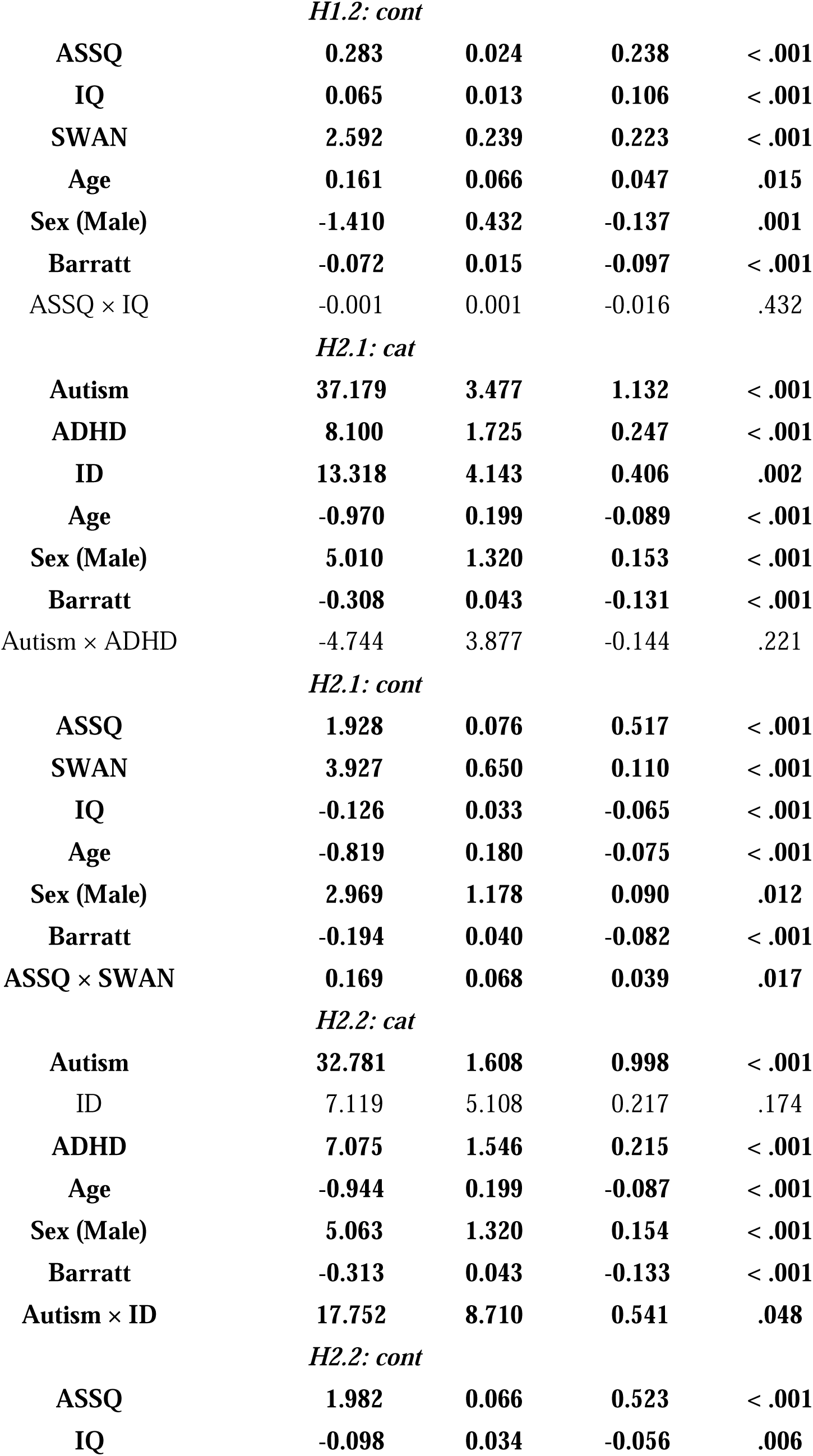
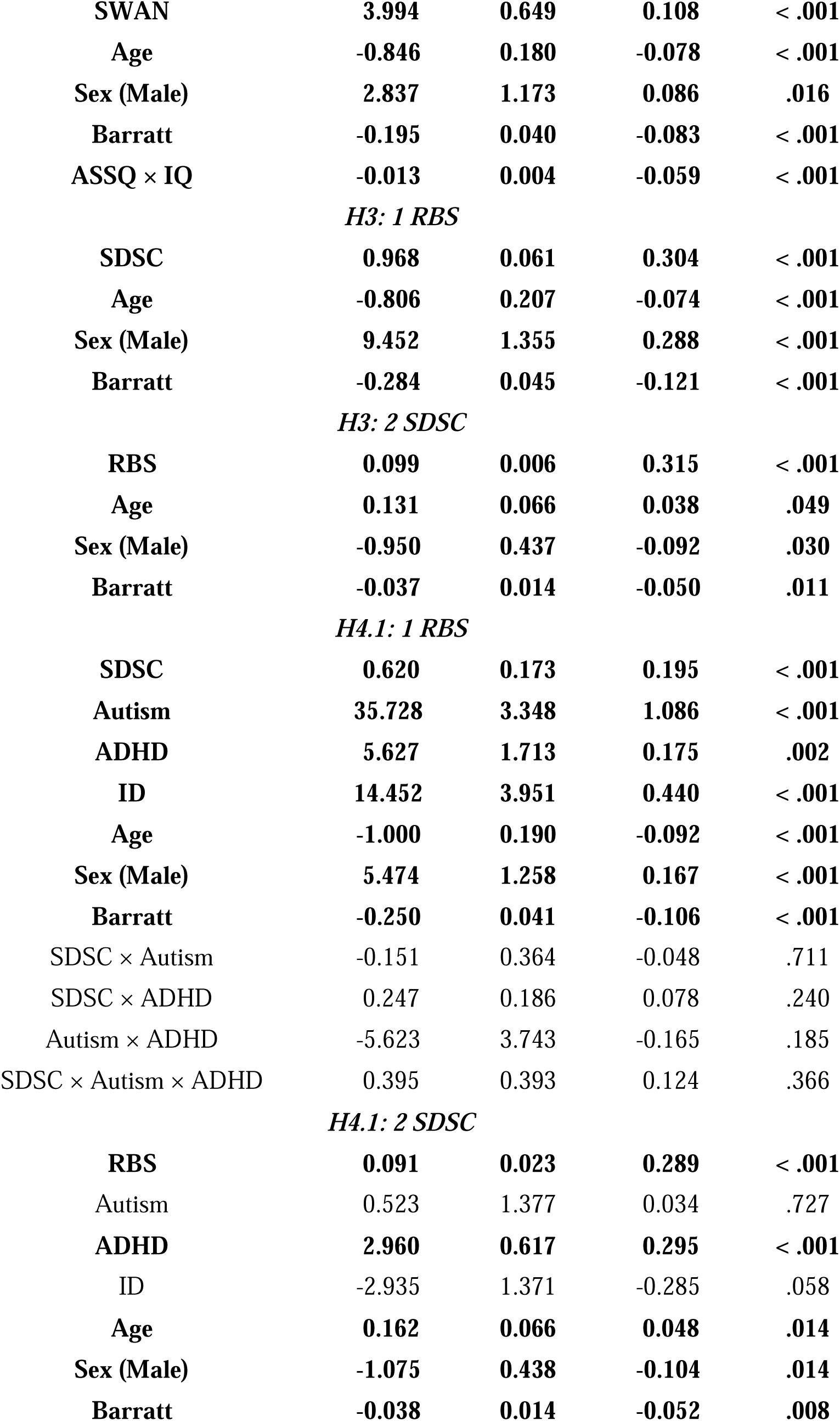
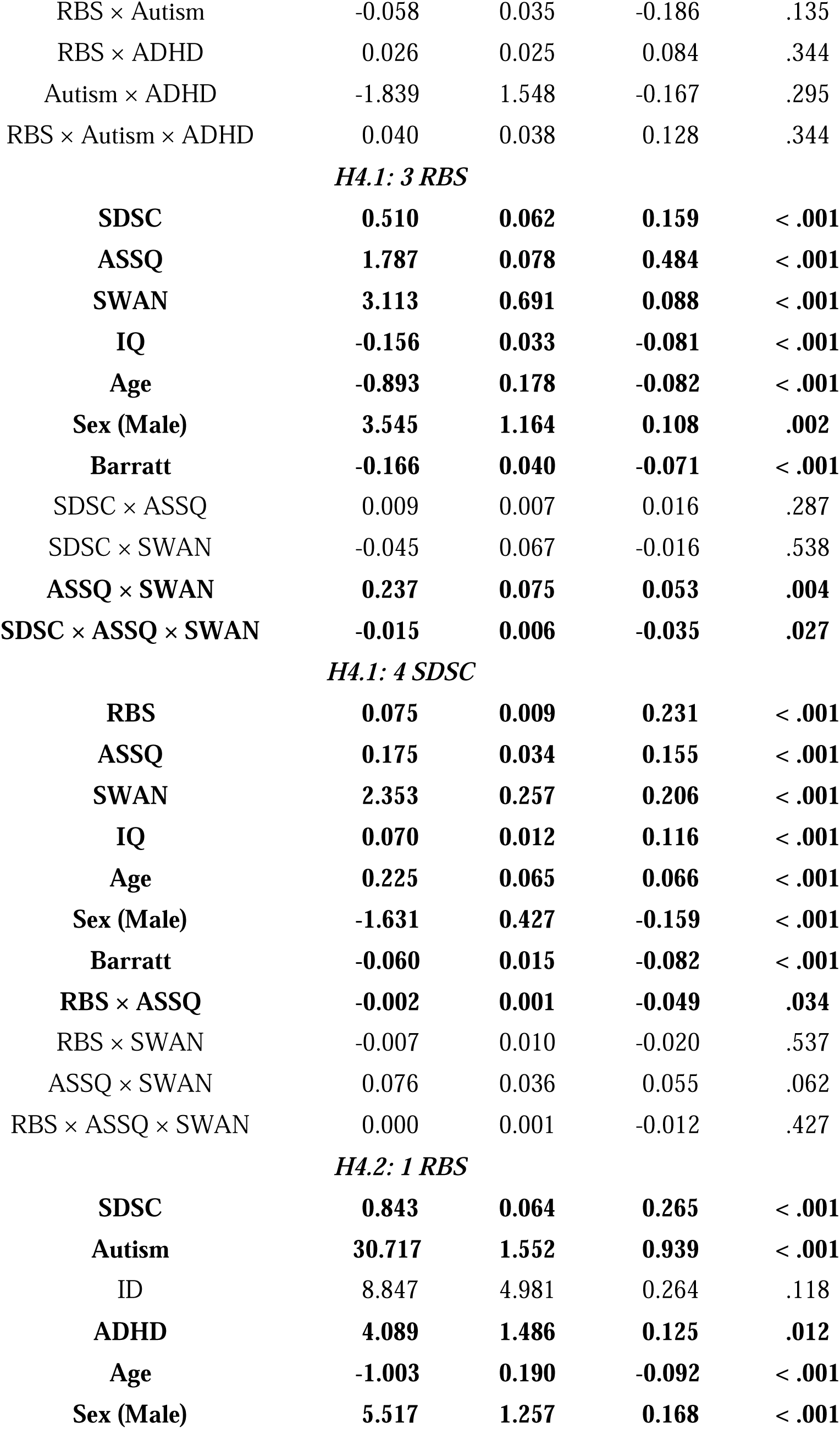
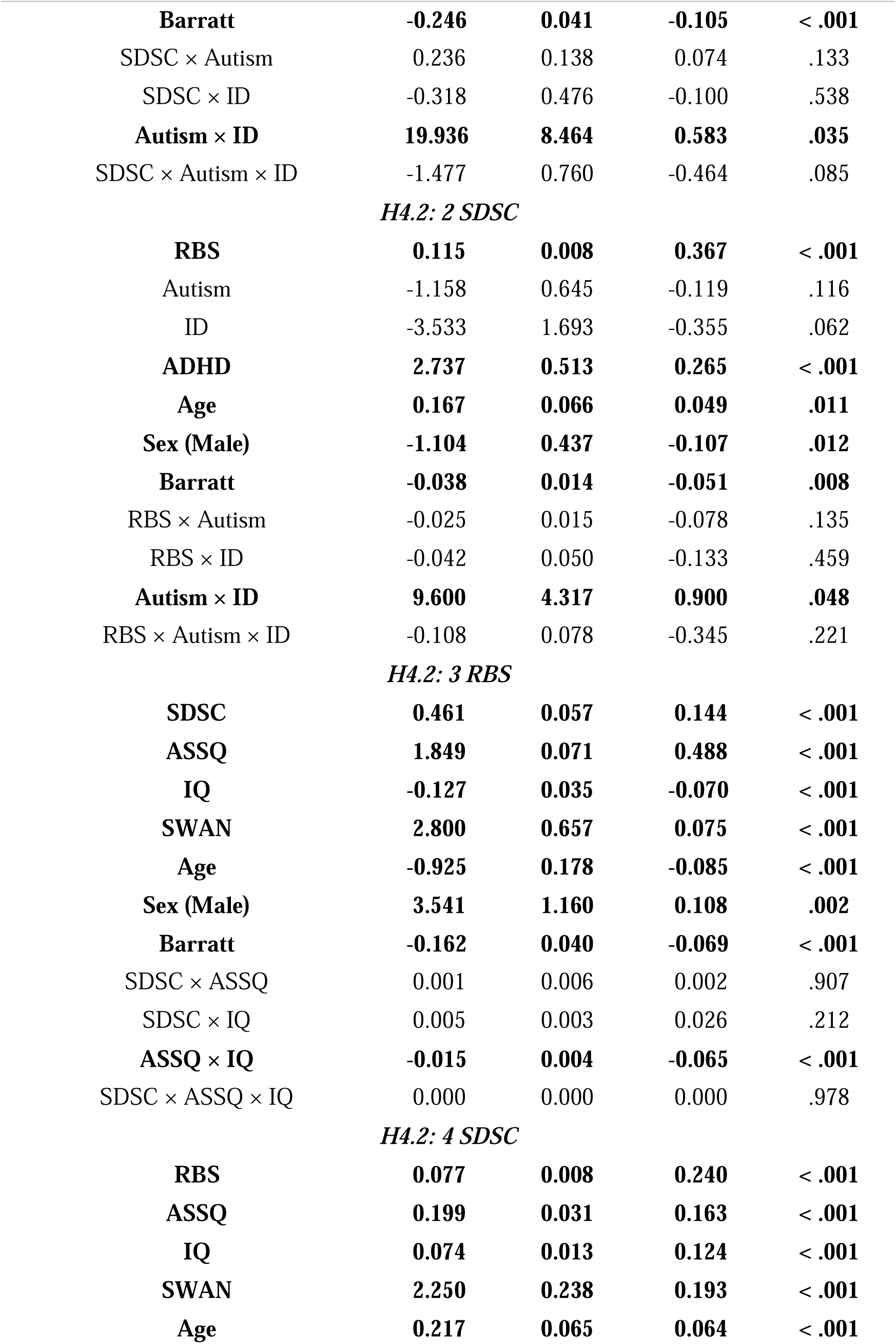
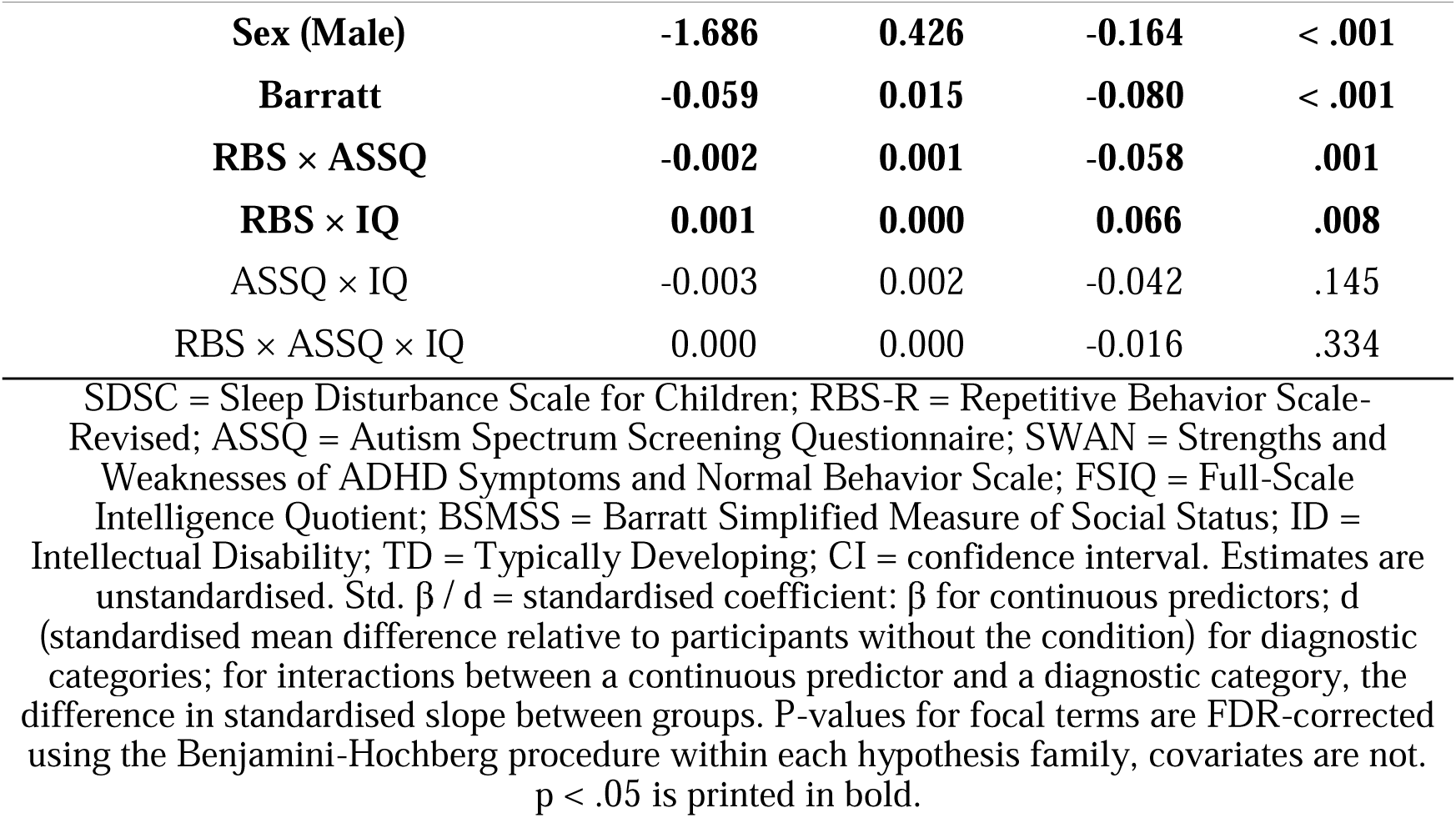
Linear regression models predicting sleep disturbance and RRBs (H1-H4).

### Exploratory analyses

RRBs were more severe in autism than in ADHD (Δ = 26.2 RBS-R points, *p* < .001) and ID (Δ = 20.2, *p* < .001), whereas ADHD and ID did not differ (*p* = .165). In all three diagnostic categories, RRBs were more severe compared to TD (all *p* ≤ .003). Higher autistic and ADHD traits as well as lower IQ were associated with more severe RRBs (all *p* < .001). This association was stronger for autistic than ADHD traits (Δβ = .43, *p* < .001) and lower IQ (Δβ = .47, *p* < .001). We found that dimensional models explained more variance than categorical models for sleep disturbance (R² = .14 vs .03; ΔR² = .11, 95% CI [.09, .13]) and RRBs (R² = .38 vs .21; ΔR² = .17, 95% CI [.13, .21]), including out of sample (cross-validated R² = .14 vs .03 and .37 vs .20, respectively). Model fits for dimensional models exceeded diagnostic models in all 8 comparisons (Vuong z = 4.00–8.23, *p* < .001, FDR-corrected). In contrast, diagnostic categories added little variance beyond traits (ΔR² ≤ .02, Supplementary Table 3). In the ADHD group, current stimulant use was associated with more sleep disturbance (d = .17, *p* = .028) but not RRB severity (d = .05, *p* = .784) compared with no psychiatric medication, and did not moderate the sleep–RRB association (*p* = .900; Supplementary Table 4).

## Discussion

Here we systematically investigated differences in sleep disturbance, RRBs, and their interaction, between ADHD, autism, ID and TD. Using both diagnostic categories and symptom dimensions, we found that differences in sleep disturbance, RRBs, and their interaction manifested mostly in symptom dimensions rather than diagnostic categories. Our results confirm previous studies^1^ which demonstrated higher rates of sleep disturbance in individuals with diagnosed ADHD relative to TD. However, autism was only associated with more sleep disturbance in one model, and ID was not linked to sleep disturbance at all. In our sample, mean SDSC scores for ADHD (M=42.2, SD=10.5), autism (M=43.1, SD=11.2), and ID (M=40.3, SD=10.9) but not TD (M=38.3, SD=8.5) exceeded the SDSC cutoff score for sleep dysfunction^33^. Considering that 40.1% of TD individuals exceed this cutoff, elevated rates of sleep disturbance in this group might explain this result (Supplementary Table 5). We also did not find evidence for our hypothesis that children with a dual diagnosis ADHD and autism or ID experience more sleep disturbance than either condition alone, which is at odds with the literature for ADHD and autism^26^. In contrast, ADHD and autistic symptom dimensions as well as IQ were positively linked to sleep disturbance. This contradicts previous findings linking more sleep problems to lower cognitive ability^41^. Since we use a community-referred sample where young people were referred for psychiatric and developmental issues, our sample is enriched for neurodevelopmental diagnoses. Individuals with high IQ are likely to exhibit at least some functional impairment to be referred for an evaluation regardless of whether they were eventually given a diagnosis. Higher IQ is also correlated with a higher preference for late sleep and wake times^42^, which is itself linked to more sleep disturbances^43^.

There was no synergistic effect of co-occurring autism and ADHD diagnoses, but high ADHD traits exacerbated the effect of autistic traits on RRBs. A combination of autistic and ADHD traits might lead to an increased need for self-soothing or sensory regulation. Co-occurring autistic and ADHD traits might also make it more difficult to suppress or mask RRBs, e.g., due to perceived judgment from observers. Concurrent high autistic traits and ADHD-linked low inhibitory control might result in freer RRB expression^44^. Both for diagnostic categories and symptoms, lower cognitive ability further reinforced the relationship between autism or autistic traits and RRB severity. This aligns with previous studies showing an association between low non-verbal IQ and RRBs in autistic children^45^. In line with previous work^18^, high RRBs and sleep disturbance were linked across the entire sample. We also found that RRBs were less strongly associated with sleep disturbance with increasing autistic traits. Possibly, individuals with more autistic traits might employ RRBs to self-regulate^46^, and therefore reduce sleep difficulties. However, in the opposite direction, sleep disturbance was less strongly associated with RRBs in individuals with high levels of both autistic and ADHD traits. Although ADHD traits were associated with more RRBs overall, ADHD and sleep disturbance might both contribute to deficits in top-down control, and thereby prevent individuals from employing sensory self-regulation strategies when autistic traits are prominent^47^.

An interesting aspect of our study is that we were largely only able to uncover differences in sleep disturbance, RRBs and their interaction based on symptom dimensions rather than diagnostic groups. Our results speak in favor of a transdiagnostic approach to assessing the relationship between RRBs and sleep. Overlaps in symptom presentation, multiple diagnoses, symptom profiles falling short of the diagnostic threshold and inconsistent diagnostic boundaries challenge research contrasting neurodevelopmental conditions^24^. Trait compositions might vary between individuals with the same diagnosis, and within the same individual depending on context and developmental stage^48^. For instance, categorical models do not distinguish between individuals with ADHD and with elevated autistic traits but no autism diagnosis, and ADHD alone. Modeling the relationship between sleep and RRBs with diagnostic categories results in a loss of variance and less statistical power relative to dimensional symptom models. Therefore, we suggest to frame the interaction of sleep issues and RRBs as a characteristic of the neurodevelopmental spectrum rather than individual conditions^2,24^.

Several limitations of our study should be acknowledged. First, the HBN dataset is cross-sectional and precludes any causal inferences about RRBs and sleep disturbance. Longitudinal studies are necessary to untangle the trajectories of both variables comparing neurodevelopmental conditions. Second, our study only used caregiver-reports and future research should integrate physiological measures, such as actigraphy or polysomnography for sleep and psychophysiological measures for sensory reactivity. Third, we defined ID using IQ and VABS scores rather than a diagnosis of intellectual developmental disorder or presence of impairment before the age of 18, which can limit the generalizability of our study to other work. However, 41 of 52 ID participants received a diagnosis of a mild (F70) or moderate (F71) intellectual disability. Fourth, the sample size description in our preregistration differs from the paper due to coding errors which assigned participants to diagnostic categories using primary diagnoses and included participants with neurological and psychiatric conditions in TD. Another limitation pertains to imbalanced diagnostic category sizes. However, they reflect greater ADHD prevalence in the population^49^. Finally, our sample was predominantly White, male and guardians were typically highly educated, potentially limiting the generalisability of our findings. Sex was recorded by caregivers and treated as a binary covariate, which precludes analyses of intersex variations.

In sum, symptom dimensions were better suited to uncover differences between sleep, RRBs, and their interaction than diagnostic categories. Our results support transdiagnostic approaches in psychiatry and encourage clinicians to assess sleep dysfunction and RRBs based on neurodevelopmental symptoms rather than diagnostic categories.

## Data and code availability

Data are available through the HBN project (https://fcon_1000.projects.nitrc.org/indi/cmi_healthy_brain_network/). Code is available here: https://osf.io/9h7cn/.

## References

1. Bruni O, Breda M, Mammarella V, Mogavero MP, Ferri R. Sleep and circadian disturbances in children with neurodevelopmental disorders. Nature Reviews Neurology. 2025;21(2):103–120. doi:10.1038/s41582-024-01052-9

2. Caserini C, Rakesh D, Lu S, et al. Developmental Pathways From Childhood Neurodevelopmental Traits to Early Adolescent Psychiatric Dimensions: The Role of Environmental and Lifestyle Factors. JAACAP open. 2026;4(2):298–309. doi:10.1016/j.jaacop.2025.11.011

3. Marten F, Keuppens L, Baeyens D, et al. Co-occurring mental health problems in adolescents with ADHD and sleep problems. Sleep Medicine. 2025;126:107–113. doi:10.1016/j.sleep.2024.12.008

4. Adamis D, Tiernan J, Langan N, Gavin B, McNicholas F. Effects of medications on sleep quality, insomnia, and circadian rhythm in adults with ADHD: A pragmatic longitudinal study. Sleep Medicine. 2026;138:108721. doi:10.1016/j.sleep.2025.108721

5. Surtees AD, Oliver C, Jones CA, Evans DL, Richards C. Sleep duration and sleep quality in people with and without intellectual disability: A meta-analysis. Sleep medicine reviews. 2018;40:135–150.

6. Van de Wouw E, Evenhuis HM, Echteld MA. Prevalence, associated factors and treatment of sleep problems in adults with intellectual disability: A systematic review. Research in developmental disabilities. 2012;33(4):1310–1332.

7. Riggins T, Ratliff EL, Horger MN, Spencer RMC. The importance of sleep for the developing brain. Current Sleep Medicine Reports. 2024;10(4):437–446. doi:10.1007/s40675-024-00307-7

8. Kamath MS, Dahm CR, Tucker JR, Huang-Pollock CL, Etter NM, Neely KA. Sensory profiles in adults with and without ADHD. Research in Developmental Disabilities. 2020;104:103696. doi:10.1016/j.ridd.2020.103696

9. Engel-Yeger B, Hardal-Nasser R, Gal E. Sensory processing dysfunctions as expressed among children with different severities of intellectual developmental disabilities. Research in Developmental Disabilities. 2011;32(5):1770–1775. doi:10.1016/j.ridd.2011.03.005

10. Márquez-Caraveo ME, Rodríguez-Valentín R, Pérez-Barrón V, et al. Children and adolescents with neurodevelopmental disorders show cognitive heterogeneity and require a person-centered approach. Scientific Reports. 2021;11(1):18463. doi:10.1038/s41598-021-97551-6

11. Bodfish JW, Symons FJ, Parker DE, Lewis MH. Varieties of Repetitive Behavior in Autism: Comparisons to Mental Retardation. Journal of Autism and Developmental Disorders. 2000;30(3):237–243. doi:10.1023/A:1005596502855

12. Brierley NJ, McDonnell CG, Parks KMA, et al. Factor Structure of Repetitive Behaviors Across Autism Spectrum Disorder and Attention-Deficit/Hyperactivity Disorder. Journal of Autism and Developmental Disorders. 2021;51(10):3391–3400. doi:10.1007/s10803-020-04800-0

13. Werkman MF, Landsman JA, Fokkens AS, et al. The Impact of the Presence of Intellectual Disabilities on Sensory Processing and Behavioral Outcomes Among Individuals with Autism Spectrum Disorders: A Systematic Review. Review Journal of Autism and Developmental Disorders. 2023;10(3):422–440. doi:10.1007/s40489-022-00301-1

14. De la Marche W, Steyaert J, Noens I. Atypical sensory processing in adolescents with an autism spectrum disorder and their non-affected siblings. Research in Autism Spectrum Disorders. 2012;6(2):639–645. doi:10.1016/j.rasd.2011.09.014

15. Roshani F, Piri R, Malek A, Michel TM, Vafaee MS. Comparison of cognitive flexibility, appropriate risk-taking and reaction time in individuals with and without adult ADHD. Psychiatry Research. 2020;284:112494. doi:10.1016/j.psychres.2019.112494

16. Lage C, Smith ES, Lawson RP. A meta-analysis of cognitive flexibility in autism spectrum disorder. Neuroscience & Biobehavioral Reviews. 2024;157:105511. doi:10.1016/j.neubiorev.2023.105511

17. Fidler DJ, Lanfranchi S. Executive function and intellectual disability: Innovations, methods and treatment. Journal of Intellectual Disability Research. 2022;66(1-2):1–8. doi:10.1111/jir.12906

18. MacDuffie KE, Munson J, Greenson J, et al. Sleep Problems and Trajectories of Restricted and Repetitive Behaviors in Children with Neurodevelopmental Disabilities. Journal of Autism and Developmental Disorders. 2020;50(11):3844–3856. doi:10.1007/s10803-020-04438-y

19. Mimouni-Bloch A, Offek H, Engel-Yeger B, et al. Association between sensory modulation and sleep difficulties in children with Attention Deficit Hyperactivity Disorder (ADHD). Sleep Medicine. 2021;84:107–113. doi:10.1016/j.sleep.2021.05.027

20. Harper L, McAnelly S, Walshe I, Ooms A, Tuffrey-Wijne IM. Behavioural sleep problems in children and adults with intellectual disabilities: An integrative literature review. Journal of Applied Research in Intellectual Disabilities. 2023;36(5):916–928. doi:10.1111/jar.13116

21. Berry K, Russell K, Frost K. Restricted and Repetitive Behaviors in Autism Spectrum Disorder: A Review of Associated Features and Presentation Across Clinical Populations. Current Developmental Disorders Reports. 2018;5(2):108–115. doi:10.1007/s40474-018-0139-0

22. Lugo J, Fadeuilhe C, Gisbert L, et al. Sleep in adults with autism spectrum disorder and attention deficit/hyperactivity disorder: A systematic review and meta-analysis. European Neuropsychopharmacology. 2020;38:1–24. doi:10.1016/j.euroneuro.2020.07.004

23. Köse S, Yılmaz H, Ocakoğlu FT, Özbaran NB. Sleep problems in children with autism spectrum disorder and intellectual disability without autism spectrum disorder. Sleep Medicine. 2017;40:69–77. doi:10.1016/j.sleep.2017.09.021

24. Michelini G, Carlisi CO, Eaton NR, et al. Where do neurodevelopmental conditions fit in transdiagnostic psychiatric frameworks? Incorporating a new neurodevelopmental spectrum. World Psychiatry. 2024;23(3):333–357. doi:10.1002/wps.21225

25. Davies LE, Oliver C. Self-injury, aggression and destruction in children with severe intellectual disability: Incidence, persistence and novel, predictive behavioural risk markers. Research in Developmental Disabilities. 2016;49–50:291-301. doi:10.1016/j.ridd.2015.12.003

26. Petti T, Gupta M, Fradkin Y, Gupta N. Management of sleep disorders in autism spectrum disorder with co-occurring attention-deficit hyperactivity disorder: Update for clinicians. BJPsych Open. 2024;10(1):e11. doi:10.1192/bjo.2023.589

27. Burbidge C, Oliver C, Moss J, et al. The association between repetitive behaviours, impulsivity and hyperactivity in people with intellectual disability. Journal of Intellectual Disability Research. 2010;54(12):1078–1092. doi:10.1111/j.1365-2788.2010.01338.x

28. Michelini G, Liao W, Lu SD, et al. The neurodevelopmental spectrum: Phenotypic architecture, etiology, predictive utility, and specificity across development. Molecular Psychiatry. Published online July 2026:1–11. doi:10.1038/s41380-026-03714-0

29. Stevens T, Peng L, Barnard-Brak L. The comorbidity of ADHD in children diagnosed with autism spectrum disorder. Research in Autism Spectrum Disorders. 2016;31:11–18. doi:10.1016/j.rasd.2016.07.003

30. Rommelse NNJ, Franke B, Geurts HM, Hartman CA, Buitelaar JK. Shared heritability of attention-deficit/hyperactivity disorder and autism spectrum disorder. European Child & Adolescent Psychiatry. 2010;19(3):281–295. doi:10.1007/s00787-010-0092-x

31. Alexander LM, Escalera J, Ai L, et al. An open resource for transdiagnostic research in pediatric mental health and learning disorders. Scientific Data. 2017;4(1):170181. doi:10.1038/sdata.2017.181

32. Townsend L, Kobak K, Kearney C, et al. Development of Three Web-Based Computerized Versions of the Kiddie Schedule for Affective Disorders and Schizophrenia Child Psychiatric Diagnostic Interview: Preliminary Validity Data. Journal of the American Academy of Child & Adolescent Psychiatry. 2020;59(2):309–325. doi:10.1016/j.jaac.2019.05.009

33. Bruni O, Ottaviano S, Guidetti V, et al. The Sleep Disturbance Scale for Children (SDSC) Construction and validation of an instrument to evaluate sleep disturbances in childhood and adolescence. Journal of Sleep Research. 1996;5(4):251–261. doi:10.1111/j.1365-2869.1996.00251.x

34. Ehlers S, Gillberg C, Wing L. A Screening Questionnaire for Asperger Syndrome and Other High-Functioning Autism Spectrum Disorders in School Age Children. Journal of Autism and Developmental Disorders. 1999;29(2):129–141. doi:10.1023/A:1023040610384

35. Swanson JM, Schuck S, Porter MM, et al. Categorical and Dimensional Definitions and Evaluations of Symptoms of ADHD: History of the SNAP and the SWAN Rating Scales. The International journal of educational and psychological assessment. 2012;10(1):51–70.

36. Wechsler D. Wechsler Adult Intelligence Scale–Fourth Edition (WAIS-IV). Published online 2008.

37. Wechsler D. Wechsler intelligence scale for children–Fifth Edition (WISC-V). Bloomington, MN: Pearson. Published online 2014.

38. Kaufman AS. Kaufman brief intelligence test. Edition 2. Published online 2004.

39. Barratt W. The Barratt simplified measure of social status measuring SES. Unpubl manuscript, Indian State Univ. Published online 2006.

40. Burger-Caplan R, Saulnier CA, Sparrow SS. Vineland Adaptive Behavior Scales. In: Encyclopedia of Clinical Neuropsychology. Springer, Cham; 2018:3597–3601. doi:10.1007/978-3-319-57111-9_1602

41. Gruber R, Laviolette R, Deluca P, Monson E, Cornish K, Carrier J. Short sleep duration is associated with poor performance on IQ measures in healthy school-age children. Sleep Medicine. 2010;11(3):289–294. doi:10.1016/j.sleep.2009.09.007

42. Ujma PP, Scherrer V. Circadian preference and intelligence - an updated meta-analysis. Chronobiology International. 2021;38(8):1215–1229. doi:10.1080/07420528.2021.1926473

43. Zhao Y, Chen H, Åkerstedt T, Zhang Z, Wang S, Tan X. Association between insomnia symptoms and chronotype—A systematic review and meta-analysis. Sleep Health. 2025;11(5):551–559. doi:10.1016/j.sleh.2025.06.011

44. Collis E, Gavin J, Russell A, Brosnan M. Autistic adults’ experience of restricted repetitive behaviours. Research in Autism Spectrum Disorders. 2022;90:101895. doi:10.1016/j.rasd.2021.101895

45. Bishop SL, Richler J, Lord C. Association Between Restricted and Repetitive Behaviors and Nonverbal IQ in Children with Autism Spectrum Disorders. Child Neuropsychology. 2006;12(4-5):247–267. doi:10.1080/09297040600630288

46. Kapp SK, Steward R, Crane L, et al. “People should be allowed to do what they like”: Autistic adults’ views and experiences of stimming. Autism. 2019;23(7):1782–1792. doi:10.1177/1362361319829628

47. D’Cruz AM, Ragozzino ME, Mosconi MW, Shrestha S, Cook EH, Sweeney JA. Reduced behavioral flexibility in autism spectrum disorders. Neuropsychology. 2013;27(2):152–160. doi:10.1037/a0031721

48. Niina A, Eyre O, Wootton R, Stergiakouli E, Thapar A, Riglin L. Exploring ADHD Symptoms and Associated Impairment across Development. Journal of Attention Disorders. 2022;26(6):822–830. doi:10.1177/10870547211025612

49. Zablotsky B, Black LI, Maenner MJ, et al. Prevalence and Trends of Developmental Disabilities among Children in the United States: 2009–2017. Pediatrics. 2019;144(4):e20190811. doi:10.1542/peds.2019-0811

