## Supplementary for "Interactions of Sleep and Repetitive Behavior in Attention-Deficit/Hyperactivity Disorder, Autism and Intellectual Disability"

Supplementary Materials

Supplementary Table 1: Clinician consensus diagnoses by diagnostic category

| ICD-10 | Diagnosis | ADHD (n = 1948) | Autism (n = 433) | ID (n = 52) | TD (n = 344) |
| --- | --- | --- | --- | --- | --- |
| F05 | Delirium due to multiple etiologies | 1 (0.1) | 0 | 0 | 0 |
| F10.20 | Alcohol Use Disorder | 1 (0.1) | 0 | 0 | 0 |
| F12.10 | Cannabis Use Disorder | 2 (0.1) | 1 (0.2) | 0 | 0 |
| F12.20 | Cannabis Use Disorder | 2 (0.1) | 1 (0.2) | 0 | 0 |
| F15.94 | Substance/Medication-Induced Bipolar and Related Disorder | 1 (0.1) | 0 | 0 | 0 |
| F17.200 | Tobacco Use Disorder | 1 (0.1) | 0 | 0 | 0 |
| F19.94 | Substance/Medication-Induced Bipolar and Related Disorder | 1 (0.1) | 1 (0.2) | 0 | 0 |
| F20.9 | Schizophrenia | 2 (0.1) | 1 (0.2) | 0 | 0 |
| F28 | Other Specified Schizophrenia Spectrum and Other Psychotic Disorder | 0 | 2 (0.5) | 0 | 0 |
| F31.81 | Bipolar II Disorder | 1 (0.1) | 0 | 0 | 0 |
| F32.0 | Major Depressive Disorder | 14 (0.7) | 1 (0.2) | 0 | 0 |
| F32.1 | Major Depressive Disorder | 14 (0.7) | 2 (0.5) | 0 | 0 |
| F32.2 | Major Depressive Disorder | 4 (0.2) | 0 | 0 | 0 |
| F32.4 | Major Depressive Disorder | 28 (1.4) | 4 (0.9) | 0 | 0 |
| F32.5 | Major Depressive Disorder | 14 (0.7) | 4 (0.9) | 0 | 0 |
| F32.8 | Other Specified Depressive Disorder | 23 (1.2) | 6 (1.4) | 0 | 0 |
| F32.9 | Unspecified Depressive Disorder | 1 (0.1) | 0 | 0 | 0 |
| F33.0 | Major Depressive Disorder | 1 (0.1) | 0 | 0 | 0 |
| F33.1 | Major Depressive Disorder | 7 (0.4) | 1 (0.2) | 0 | 0 |
| F33.2 | Major Depressive Disorder | 4 (0.2) | 2 (0.5) | 0 | 0 |
| F33.3 | Major Depressive Disorder | 1 (0.1) | 0 | 0 | 0 |
| F33.41 | Major Depressive Disorder | 11 (0.6) | 0 | 0 | 0 |
| F33.42 | Major Depressive Disorder | 5 (0.3) | 2 (0.5) | 0 | 0 |
| F34.0 | Cyclothymic Disorder | 1 (0.1) | 0 | 0 | 0 |
| F34.1 | Persistent Depressive Disorder (Dysthymia) | 33 (1.7) | 5 (1.2) | 1 (1.9) | 0 |
| F34.8 | Disruptive Mood Dysregulation Disorder | 20 (1.0) | 3 (0.7) | 0 | 0 |
| F40.0 | Agoraphobia | 6 (0.3) | 2 (0.5) | 0 | 0 |
| F40.10 | Social Anxiety (Social Phobia) | 195 (10.0) | 34 (7.9) | 3 (5.8) | 0 |
| F40.218 | Specific Phobia | 51 (2.6) | 20 (4.6) | 3 (5.8) | 0 |
| F40.228 | Specific Phobia | 12 (0.6) | 2 (0.5) | 0 | 0 |
| F40.230 | Specific Phobia | 5 (0.3) | 2 (0.5) | 0 | 0 |
| F40.231 | Specific Phobia | 65 (3.3) | 21 (4.8) | 1 (1.9) | 0 |
| F40.232 | Specific Phobia | 4 (0.2) | 1 (0.2) | 1 (1.9) | 0 |
| F40.233 | Specific Phobia | 2 (0.1) | 0 | 0 | 0 |
| F40.248 | Specific Phobia | 64 (3.3) | 15 (3.5) | 4 (7.7) | 0 |
| F40.298 | Specific Phobia | 13 (0.7) | 7 (1.6) | 2 (3.8) | 0 |
| F41.0 | Panic Disorder | 5 (0.3) | 2 (0.5) | 0 | 0 |
| F41.1 | Generalized Anxiety Disorder | 346 (17.8) | 78 (18.0) | 5 (9.6) | 0 |
| F41.8 | Other Specified Anxiety Disorder | 87 (4.5) | 16 (3.7) | 2 (3.8) | 0 |
| F41.9 | Unspecified Anxiety Disorder | 3 (0.2) | 0 | 0 | 0 |
| F42 | Obsessive-Compulsive Disorder | 62 (3.2) | 24 (5.5) | 0 | 0 |
| F43.10 | Posttraumatic Stress Disorder | 28 (1.4) | 6 (1.4) | 1 (1.9) | 0 |
| F43.21 | Adjustment Disorders | 8 (0.4) | 0 | 0 | 0 |
| F43.22 | Adjustment Disorders | 9 (0.5) | 1 (0.2) | 0 | 0 |
| F43.23 | Adjustment Disorders | 10 (0.5) | 4 (0.9) | 0 | 0 |
| F43.24 | Adjustment Disorders | 2 (0.1) | 0 | 0 | 0 |
| F43.25 | Adjustment Disorders | 15 (0.8) | 2 (0.5) | 0 | 0 |
| F43.8 | Other Specified Trauma- and Stressor-Related Disorder | 14 (0.7) | 6 (1.4) | 2 (3.8) | 0 |
| F43.9 | Unspecified Trauma- and Stressor-Related Disorder | 2 (0.1) | 0 | 0 | 0 |
| F45.21 | Illness Anxiety Disorder | 1 (0.1) | 0 | 0 | 0 |
| F45.22 | Body Dysmorphic Disorder | 1 (0.1) | 0 | 0 | 0 |
| F50.8 | Other Specified Feeding or Eating Disorder | 6 (0.3) | 3 (0.7) | 0 | 0 |
| F51.3 | Non-Rapid Eye Movement Sleep Arousal Disorders | 1 (0.1) | 0 | 0 | 0 |
| F51.4 | Non-Rapid Eye Movement Sleep Arousal Disorders | 2 (0.1) | 1 (0.2) | 0 | 0 |
| F63.3 | Trichotillomania (Hair-Pulling Disorder) | 8 (0.4) | 4 (0.9) | 2 (3.8) | 0 |
| F63.81 | Intermittent Explosive Disorder | 9 (0.5) | 0 | 0 | 0 |
| F64.1 | Gender Dysphoria in Adolescents and Adults | 2 (0.1) | 0 | 0 | 0 |
| F64.2 | Gender Dysphoria in Children | 1 (0.1) | 0 | 0 | 0 |
| F70 | Intellectual Disability-Mild | 38 (2.0) | 16 (3.7) | 35 (67.3) | 0 |
| F71 | Intellectual Disability-Moderate | 5 (0.3) | 2 (0.5) | 6 (11.5) | 0 |
| F80.0 | Speech Sound Disorder | 89 (4.6) | 25 (5.8) | 6 (11.5) | 0 |
| F80.81 | Child Onset Fluency Disorder (Stuttering) | 12 (0.6) | 2 (0.5) | 0 | 0 |
| F80.89 | Social (Pragmatic) Communication Disorder | 18 (0.9) | 2 (0.5) | 0 | 0 |
| F80.9 | Language Disorder | 272 (14.0) | 74 (17.1) | 12 (23.1) | 0 |
| F81.0 | Specific Learning Disorder with Impairment in Reading | 377 (19.4) | 58 (13.4) | 12 (23.1) | 0 |
| F81.2 | Specific Learning Disorder with Impairment in Mathematics | 205 (10.5) | 41 (9.5) | 9 (17.3) | 0 |
| F81.81 | Specific Learning Disorder with Impairment in Written Expression | 172 (8.8) | 32 (7.4) | 5 (9.6) | 0 |
| F82 | Developmental Coordination Disorder | 32 (1.6) | 15 (3.5) | 2 (3.8) | 0 |
| F84.0 | Autism Spectrum Disorder | 343 (17.6) | 433 (100.0) | 18 (34.6) | 0 |
| F88 | Other Specified Neurodevelopmental Disorder | 2 (0.1) | 0 | 0 | 0 |
| F89 | Unspecified Neurodevelopmental Disorder | 1 (0.1) | 0 | 0 | 0 |
| F90.0 | ADHD-Inattentive Type | 861 (44.2) | 130 (30.0) | 18 (34.6) | 0 |
| F90.1 | ADHD-Hyperactive/Impulsive Type | 107 (5.5) | 10 (2.3) | 0 | 0 |
| F90.2 | ADHD-Combined Type | 853 (43.8) | 191 (44.1) | 17 (32.7) | 0 |
| F90.8 | Other Specified Attention-Deficit/Hyperactivity Disorder | 125 (6.4) | 12 (2.8) | 1 (1.9) | 0 |
| F90.9 | Unspecified Attention-Deficit/Hyperactivity Disorder | 2 (0.1) | 0 | 0 | 0 |
| F91.1 | Conduct Disorder-Childhood-onset type | 3 (0.2) | 1 (0.2) | 0 | 0 |
| F91.2 | Conduct Disorder-Adolescent-onset type | 5 (0.3) | 0 | 0 | 0 |
| F91.3 | Oppositional Defiant Disorder | 358 (18.4) | 56 (12.9) | 4 (7.7) | 0 |
| F91.8 | Other Specified Disruptive, Impulse-Control, and Conduct Disorder | 6 (0.3) | 2 (0.5) | 0 | 0 |
| F91.9 | Conduct Disorder-Unspecified onset | 1 (0.1) | 0 | 0 | 0 |
| F93.0 | Separation Anxiety | 137 (7.0) | 27 (6.2) | 0 | 0 |
| F94.0 | Selective Mutism | 11 (0.6) | 0 | 0 | 0 |
| F94.1 | Reactive Attachment Disorder | 3 (0.2) | 0 | 1 (1.9) | 0 |
| F94.2 | Disinhibited Social Engagement Disorder | 1 (0.1) | 0 | 0 | 0 |
| F95.0 | Provisional Tic Disorder | 41 (2.1) | 7 (1.6) | 0 | 0 |
| F95.1 | Persistent (Chronic) Motor or Vocal Tic Disorder | 86 (4.4) | 24 (5.5) | 3 (5.8) | 0 |
| F95.2 | Tourettes Disorder | 39 (2.0) | 15 (3.5) | 0 | 0 |
| F95.8 | Other Specified Tic Disorder | 9 (0.5) | 4 (0.9) | 0 | 0 |
| F95.9 | Unspecified Tic Disorder | 2 (0.1) | 0 | 0 | 0 |
| F98.0 | Enuresis | 213 (10.9) | 53 (12.2) | 8 (15.4) | 0 |
| F98.1 | Encopresis | 57 (2.9) | 17 (3.9) | 0 | 0 |
| F98.3 | Pica in Children | 3 (0.2) | 1 (0.2) | 0 | 0 |
| G47.00 | Insomnia Disorder | 6 (0.3) | 3 (0.7) | 0 | 0 |
| L98.1 | Excoriation (Skin-Picking) Disorder | 27 (1.4) | 7 (1.6) | 2 (3.8) | 2 (0.6) |
| N39.498 | Other Specified Elimination Disorder with Urinary Symptoms | 1 (0.1) | 0 | 0 | 0 |
| No ICD 10 Code | Specific Phobia | 23 (1.2) | 2 (0.5) | 0 | 18 (5.2) |
| R15.9 | Other Specified Elimination Disorder with Fecal Symptoms | 2 (0.1) | 1 (0.2) | 0 | 0 |
| R41.83 | Borderline Intellectual Functioning | 13 (0.7) | 7 (1.6) | 1 (1.9) | 2 (0.6) |
| Z72.0 | Tobacco Use Disorder | 3 (0.2) | 0 | 0 | 0 |
| Values represent the sample size (% of diagnostic category). Diagnostic categories overlap, so columns do not sum to the total sample size. | | | | | |
| SDSC = Sleep Disturbance Scale for Children; RBS-R = Repetitive Behavior Scale-Revised; ASSQ = Autism Spectrum Screening Questionnaire; SWAN = Strengths and Weaknesses of ADHD Symptoms and Normal Behavior Scale; FSIQ = Full-Scale Intelligence Quotient; BSMSS = Barratt Simplified Measure of Social Status; ID = Intellectual Disability; TD = Typically Developing; CI = confidence interval. | | | | | |

Supplementary Table 2: Questionnaire measures

| Construct | Instrument | Items | Rating scale | Structure | Score analyzed |
| --- | --- | --- | --- | --- | --- |
| Sleep disturbance | Sleep Disturbance Scale for Children (SDSC; Bruni et al., 1996) | 26 | 1 (never) to 5 (always) | Subscales: initiating/maintaining sleep, sleep breathing disorders, arousal, sleep–wake transition, excessive somnolence, hyperhidrosis | Total score (26–130) |
| Restricted and repetitive behavior | Repetitive Behavior Scale–Revised (RBS-R; Bodfish et al., 2000) | 43 | 0 (behavior does not occur) to 3 (behavior occurs and is a severe problem); final item 0–100 | Subscales: ritualistic/sameness, stereotypic, self-injurious, compulsive, restricted interests | Total score (0–129) |
| Autistic traits | Autism Spectrum Screening Questionnaire (ASSQ; Ehlers et al., 1999) | 27 | 0 (no) to 2 (yes) |  | Total score (0–54) |
| ADHD traits | Strengths and Weaknesses of ADHD Symptoms and Normal Behavior Scale (SWAN; Swanson et al., 2012) | 18 | −3 (far below average) to +3 (far above average) | Assessment relative to peers over the past month | Mean item score |
| Cognitive ability | WAIS-IV (>17 y; Wechsler, 2008); WISC-V (6–17 y; Wechsler, 2014); KBIT-2 (<6 y; Kaufman, 2004) | 15 subtests 16 subtests 3 subtests | Direct standardized assessment | Clinician-administered | Full-scale IQ |
| Adaptive behavior | Vineland Adaptive Behavior Scales, Third Edition, Parent/Caregiver Rating Form (Sparrow et al., 2016) | 502 | 0 (never performed without help) to 2 (usually occurs without help) | Caregiver-completed | Standard scores (M = 100, SD = 15) |
| Socioeconomic status | Barratt Simplified Measure of Social Status (BSMSS; Barratt, 2006) | 4 | Education: 3 (< 7th grade) to 21 (graduate degree); Occupation: 5 (low status) to 45 (high status) |  | Total score (8–66) |
| Questionnaires were completed by caregivers unless stated otherwise. ADHD = attention-deficit/hyperactivity disorder; ID = intellectual disability. | | | | | |

Supplementary Table 3: Comparison of diagnostic category and dimensional symptom models.

| Model | N | R² cat | R² dim | ΔAIC | cvR² cat | cvR² dim | Adding dimensional symptoms | Adding diagnostic categories | ΔR² dim − cat [95% CI] | Vuong z |
| --- | --- | --- | --- | --- | --- | --- | --- | --- | --- | --- |
| H1.1: SDSC | 2,376 | 0.035 | 0.143 | -283.9 | 0.028 | 0.136 | F(4, 2364) = 81.21, ΔR² = 0.117, p < .001 | F(4, 2364) = 5.50, ΔR² = 0.008, p < .001 | 0.109 [0.086, 0.133] | 8.19, p < .001 |
| H1.2: SDSC | 2,376 | 0.035 | 0.143 | -283.8 | 0.027 | 0.137 | F(4, 2364) = 81.65, ΔR² = 0.117, p < .001 | F(4, 2364) = 5.91, ΔR² = 0.008, p < .001 | 0.109 [0.084, 0.133] | 8.23, p < .001 |
| H2.1: RBS-R | 2,376 | 0.205 | 0.377 | -580.3 | 0.198 | 0.372 | F(4, 2364) = 182.62, ΔR² = 0.188, p < .001 | F(4, 2364) = 14.98, ΔR² = 0.015, p < .001 | 0.172 [0.133, 0.211] | 7.97, p < .001 |
| H2.2: RBS-R | 2,376 | 0.206 | 0.380 | -586.7 | 0.199 | 0.374 | F(4, 2364) = 184.45, ΔR² = 0.189, p < .001 | F(4, 2364) = 14.77, ΔR² = 0.015, p < .001 | 0.174 [0.132, 0.212] | 7.99, p < .001 |
| H4.1: SDSC → RBS-R | 2,376 | 0.280 | 0.397 | -420.2 | 0.269 | 0.387 | F(7, 2357) = 78.88, ΔR² = 0.137, p < .001 | F(7, 2357) = 11.52, ΔR² = 0.020, p < .001 | 0.117 [0.081, 0.152] | 6.00, p < .001 |
| H4.1: RBS-R → SDSC | 2,376 | 0.127 | 0.173 | -126.8 | 0.117 | 0.162 | F(7, 2357) = 25.29, ΔR² = 0.061, p < .001 | F(7, 2357) = 6.48, ΔR² = 0.016, p < .001 | 0.045 [0.025, 0.068] | 4.00, p < .001 |
| H4.2: SDSC → RBS-R | 2,376 | 0.282 | 0.397 | -414.7 | 0.270 | 0.387 | F(7, 2357) = 79.37, ΔR² = 0.137, p < .001 | F(7, 2357) = 12.73, ΔR² = 0.022, p < .001 | 0.115 [0.077, 0.151] | 5.82, p < .001 |
| H4.2: RBS-R → SDSC | 2,376 | 0.128 | 0.175 | -130.5 | 0.116 | 0.164 | F(7, 2357) = 25.01, ΔR² = 0.060, p < .001 | F(7, 2357) = 5.67, ΔR² = 0.014, p < .001 | 0.047 [0.027, 0.068] | 4.34, p < .001 |
| Each row compares the categorical (cat) with the dimensional model (dim) fitted to the same participants and adjusted for age, sex and BSMSS. ΔAIC = dimensional minus categorical; negative values favor the dimensional model. cvR² = mean out-of-sample R² across 50 repetitions of 10-fold cross-validation with identical folds for both models. Adding = nested F-test of each set of predictors added to a model containing the other set; p-values are FDR-corrected within column. ΔR² dim − cat = difference in R² between dimensional and categorical models with 95% percentile bootstrap CI (2,000 resamples). Vuong z = non-nested likelihood-ratio test; positive values favor the dimensional model; p-values FDR-corrected. Exploratory analysis. | | | | | | | | | | |
| SDSC = Sleep Disturbance Scale for Children; RBS-R = Repetitive Behavior Scale-Revised; ASSQ = Autism Spectrum Screening Questionnaire; SWAN = Strengths and Weaknesses of ADHD Symptoms and Normal Behavior Scale; FSIQ = Full-Scale Intelligence Quotient; BSMSS = Barratt Simplified Measure of Social Status; ID = Intellectual Disability; TD = Typically Developing; CI = confidence interval. | | | | | | | | | | |

Supplementary Table 4: Medication status predicts sleep disturbance but not RRBs or their association in participants with ADHD.

| Predictor | Estimate | SE | Std. β | t | p |
| --- | --- | --- | --- | --- | --- |
| ***Medication status: SDSC*** | | | | | |
| Other psychiatric medication (vs. no psychiatric medication) | 1.777 | 2.130 | 0.168 | 0.83 | .784 |
| Non-stimulant (vs. no psychiatric medication) | -0.413 | 1.952 | -0.039 | -0.21 | .900 |
| Stimulant (vs. no psychiatric medication) | 1.752 | 0.639 | 0.166 | 2.74 | .028 |
| Age | 0.094 | 0.092 | 0.026 | 1.02 | .308ᵃ |
| Sex (male) | -0.946 | 0.539 | -0.089 | -1.75 | .079ᵃ |
| BSMSS | -0.078 | 0.018 | -0.101 | -4.42 | < .001ᵃ |
| SWAN | 3.145 | 0.324 | 0.233 | 9.70 | < .001ᵃ |
| ***Medication status: RBS-R*** | | | | | |
| Other psychiatric medication (vs. no psychiatric medication) | 23.668 | 6.475 | 0.715 | 3.66 | .002 |
| Non-stimulant (vs. no psychiatric medication) | 1.125 | 5.935 | 0.034 | 0.19 | .900 |
| Stimulant (vs. no psychiatric medication) | 1.515 | 1.943 | 0.046 | 0.78 | .784 |
| Age | -0.505 | 0.279 | -0.044 | -1.81 | .071ᵃ |
| Sex (male) | 6.926 | 1.639 | 0.209 | 4.23 | < .001ᵃ |
| BSMSS | -0.345 | 0.053 | -0.144 | -6.47 | < .001ᵃ |
| SWAN | 11.718 | 0.986 | 0.277 | 11.89 | < .001ᵃ |
| ***SDSC × Medication: RBS-R*** | | | | | |
| SDSC (centered) | 0.880 | 0.081 | 0.281 | 10.87 | < .001ᵃ |
| Other psychiatric medication (vs. no psychiatric medication) | 25.118 | 6.324 | 0.760 | 3.97 | .001 |
| Non-stimulant (vs. no psychiatric medication) | 2.070 | 5.718 | 0.062 | 0.36 | .900 |
| Stimulant (vs. no psychiatric medication) | 0.104 | 1.881 | 0.003 | 0.06 | .956 |
| Age | -0.580 | 0.268 | -0.051 | -2.16 | .031ᵃ |
| Sex (male) | 7.705 | 1.574 | 0.233 | 4.89 | < .001ᵃ |
| BSMSS | -0.275 | 0.052 | -0.114 | -5.34 | < .001ᵃ |
| SWAN | 9.045 | 0.971 | 0.214 | 9.31 | < .001ᵃ |
| SDSC (centered) × Other psychiatric medication (vs. no psychiatric medication) | -1.537 | 0.609 | -0.491 | -2.52 | .035 |
| SDSC (centered) × Non-stimulant (vs. no psychiatric medication) | 0.723 | 0.652 | 0.231 | 1.11 | .603 |
| SDSC (centered) × Stimulant (vs. no psychiatric medication) | -0.077 | 0.154 | -0.025 | -0.50 | .900 |
| ***RBS-R × Medication: SDSC*** | | | | | |
| RBS-R (centered) | 0.093 | 0.009 | 0.292 | 10.80 | < .001ᵃ |
| Other psychiatric medication (vs. no psychiatric medication) | 2.693 | 2.265 | 0.255 | 1.19 | .603 |
| Non-stimulant (vs. no psychiatric medication) | -0.533 | 1.872 | -0.050 | -0.28 | .900 |
| Stimulant (vs. no psychiatric medication) | 1.583 | 0.613 | 0.150 | 2.58 | .035 |
| Age | 0.140 | 0.088 | 0.038 | 1.58 | .114ᵃ |
| Sex (male) | -1.536 | 0.520 | -0.145 | -2.96 | .003ᵃ |
| BSMSS | -0.045 | 0.017 | -0.059 | -2.64 | .008ᵃ |
| SWAN | 2.072 | 0.323 | 0.153 | 6.41 | < .001ᵃ |
| RBS-R (centered) × Other psychiatric medication (vs. no psychiatric medication) | -0.148 | 0.047 | -0.464 | -3.14 | .010 |
| RBS-R (centered) × Non-stimulant (vs. no psychiatric medication) | 0.014 | 0.060 | 0.045 | 0.24 | .900 |
| RBS-R (centered) × Stimulant (vs. no psychiatric medication) | 0.007 | 0.017 | 0.022 | 0.42 | .900 |
| Exploratory analysis, not preregistered. Linear regression models in participants with ADHD (n = 1,783), adjusted for age, sex, BSMSS and ADHD traits (SWAN). Medication status was based on caregiver-reported current psychiatric medication at intake (treatment history instrument): no psychiatric medication (n = 1,341), other psychiatric medication (n = 24), non-stimulant (n = 28), stimulant (n = 390). Participants taking only non-psychiatric medications were grouped with those taking no medication. | | | | | |
| P-values for medication terms are FDR-corrected (Benjamini-Hochberg) across all models. | | | | | |
| ᵃ Not a medication term; p-value uncorrected. | | | | | |
| Std. β for medication groups = standardized mean difference (d). | | | | | |
| SDSC and RBS-R were mean-centered in interaction models. | | | | | |

Supplementary Table 5: Presence of sleep disturbance in each diagnostic category based on a cut-off SDSC total score of 39.

| Diagnostic category | N | SDSC, M (SD) | Above cutoff, n (%) | Below cutoff, n (%) |
| --- | --- | --- | --- | --- |
| ADHD | 1,948 | 42.2 (10.5) | 1118 (57.4) | 830 (42.6) |
| Autism | 433 | 43.1 (11.2) | 261 (60.3) | 172 (39.7) |
| ID | 52 | 40.3 (10.9) | 27 (51.9) | 25 (48.1) |
| TD | 344 | 38.3 (8.5) | 138 (40.1) | 206 (59.9) |
| SDSC total score ≥ 39 flags sleep disturbance. Diagnostic categories overlap so that rows do not sum to the total sample size. | | | | |
| SDSC = Sleep Disturbance Scale for Children; RBS-R = Repetitive Behavior Scale-Revised; ASSQ = Autism Spectrum Screening Questionnaire; SWAN = Strengths and Weaknesses of ADHD Symptoms and Normal Behavior Scale; FSIQ = Full-Scale Intelligence Quotient; BSMSS = Barratt Simplified Measure of Social Status; ID = Intellectual Disability; TD = Typically Developing; CI = confidence interval. | | | | |
